# Time grid-based isomer specific N-glycan analysis and detection of bisecting Lewis X in human brain

**DOI:** 10.1101/2021.04.14.439640

**Authors:** Johannes Helm, Clemens Gruber, Andreas Thader, Jonathan Urteil, Johannes Führer, David Stenitzer, Daniel Maresch, Laura Neumann, Martin Pabst, Friedrich Altmann

## Abstract

The importance of protein glycosylation in the biomedical field demands for methods capable of resolving and identifying isomeric structures of N-glycans. However, the unambiguous identification of isomeric structures from complex mixtures is currently not reasonably realized even by the most sophisticated approaches. Here we present a novel approach which uses stable isotope labelled reference N-glycans to establish a retention time grid (glyco-TiGr) on porous graphitized carbon. This furthermore enables retention as the primary criterion for the structural assignment of isomeric N-glycans.

Moreover, we biosynthesized forty natural isomers of the fundamental N-glycan type consisting of five hexoses, four N-acetylhexosamines and one fucose residue. Nearly all of these isomers occupied unique positions on the retention time grid. Reference glycan assisted retention time determination with deci-minute accuracy narrowed the assignment space to very few, often only one possible glycan isomer.

Application of the glyco-TiGr approach revealed yet undescribed isomers of Lewis x determinants in multimeric human IgA and hybrid type N-glycans in human brain with galactose and even fucose linked to the bisecting N-acetylglucosamine. Thus, the brain N-glycome displayed a degree of sophistication commensurate with this organ’s role.

## INTRODUCTION

Individual glycoforms of glycoproteins confer biological properties and are increasingly considered as markers for health and disease^1–5^. The introduction of fluorescent labels for liquid chromatography or capillary electrophoresis ^6–10^, the development of ESI-MS ^11^ and MALDI-TOF MS for native or derivatized glycans ^12^ and the versatility of LC-ESI-MS of glycopeptides ^13, 14^ have brought glycan analysis into the “omics” era. The price for high throughput, however, all too often is an oversimplification of results by ignoring the possible, often simultaneous occurrence of a large number of isomers. Identification of structural isoforms of N-glycans is still one of the ‘grand challenges’ in glycomics of complex ^15, 16^, which arguably requires separation of isomers ^17^. HILIC-HPLC with amide-functionalized stationary phases has found wide applications offering the advantage of identical molar response of all N-glycan species in a sample. Correct quantitation requires good separation of peaks, which is facilitated by ultra-high-performance HILIC columns ^8, 18^, which, however reach limits with triantennary glycans. Retention is based on number and - to a limited degree - on position of the hydroxyl groups of a glycan. Thus, HILIC-HPLC can just about distinguish between core and outer arm fucosylation ^19^. However, the retention time differences provided even by UPLC-HILIC columns can hardly be seen as sufficient to discriminate the over 40 isomers of the fundamental N-glycan composition of five hexoses, four N-acetyl-hexosamines and one fucose (H5N4F1) (Figure 1). This is regrettable given the outstanding diversity and significance of fucose containing structural elements ^20^. The recently introduced coupling of HILIC-HPLC and ESI-MS certainly improves the cognitive gain ^21^, but does not substantially increase the ability to separate and annotate N-glycan isomers.

**Fig. 1.**
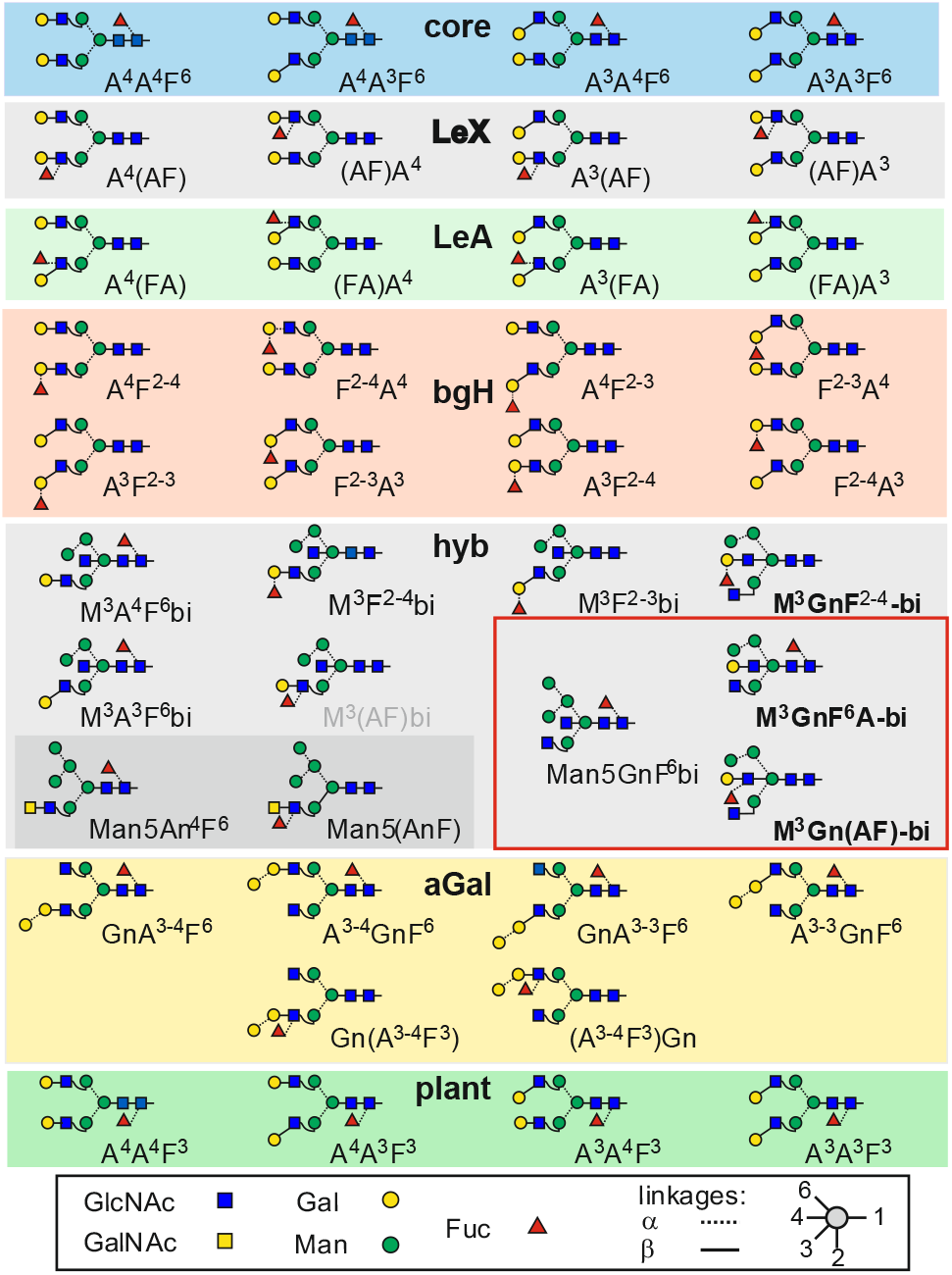
Fundamental N-glycan structures composed of five hexoses, four *N*-acetylhexosamines and one fucose (H5N4F1) used in this study. The red box comprises brain glycan structures. Names of supposedly novel structures are boldfaced. The abbreviation system is explained in the Supplementary Information.

A much higher resolution of isomeric glycans is achieved by porous graphitized carbon chromatography (PGC), which exert their superb ability only with underivatized glycans and hence *a priori* require MS detection, which is often performed in negative ion mode to exploit the information content of MS/MS spectra ^22–26^.Though currently being the most promising approach for in-depth glycome analysis, PGC-LC-MS poses the need to tackle the notoriously somewhat unstable elution times in PGC chromatography. This problem could be met by isotope labelled internal standards. A meagre 1 Da difference can be introduced by performing the required reduction step with NaBD_4_, but this cannot shift the labelled glycans away from the isotope pattern of the non-labelled analyte. Chemical synthesis has recently been employed to synthesize a range of ^13^C-labelled N-glycans ^27^. Such internal standards allow correct quantification of glycans from IgG or serum and can also be applied for PGC-LC-MS.

Retention times for a number of N-glycans can be found at https://www.glycostore.org/showPgc ^22^. However, an elution time matching that of a certain glycan in this data base or coinciding with an internal standard does not rule out the possibility of this glycan being another isomer with incidentally the same retention time. This uncertainty remains, unless the incidental co-elution of another isomer can be ruled out either by negative mode MS/MS fragment spectra ^23–25^ or by knowing the retention times of all possible isomers. In the absence of such knowledge, a recent highly comprehensive study of brain glycans admittedly had to refrain from assigning structures to many of the glycan peaks ^23^. This and other studies revealed an unusually high amount of bisecting GlcNAc in brains of various species ^23, 28, 29^.

As of today, probably all human and mammalian glycosyltransferases and structural features are known and thus the glycome space, *i.e.* the entirety of all N-glycan structures, in particular all possible isomers of a given mass level can be predicted. Establishing retention times of all possible isomers relative to each other by a shape-selective separation mechanism, which is maximally realized by PGC-LC, provides a rational approach for structural assignment by ruling out all isomers not fulfilling the retention criterium. First steps in this latter direction have been attempted with a selection of the most likely occurring disialo N-glycans ^30^ and with oligomannosidic N-glycans ^31^.

In the present work, we extend the range of synthetic reference structures by complex-type N-glycans with fucose and introduce a solution for overcoming retention time differences, which may derive from different columns, gradients, operators or frequently observed ‘aging’ (redox reactions) of the porous graphitized carbon separation phase. Thirty-six isomers of glycans containing five hexoses, four *N*-acetylhexosamines and one fucose (H5N4F1) that may occur in mammals were bio-synthetized, whereby highly un-likely arrangements and blood group determinants were spared out. In addition, four isomers of relevance in plant biotechnology were provided. To prepare for the obfuscating effects of retention time instability, the PGC runs were conducted in the presence of eight synthetic (internal) standards with characteristic mass labels. With the help of this glycan retention <u>Ti</u>me <u>Gr</u>id (glyco-TiGr), individual chromatographic runs – some with strongly deviating conditions - could be projected to a “mother” chromatogram with deci-minute precision. Experimental retention times are thereby converted to “virtual minutes” (vi-min) that are used much in the sense of the well-tried “glucose units”, which are known from hydrophilic interaction chromatography ^8, 9^ and capillary gel electro-phoresis ^32, 33^. The obtained retention time and reference glycan library was applied to the analysis of multimeric human IgA and brains from mouse, pig and human.

## RESULTS

To fathom the potential of PGC for distinguishing the large number of glycans with the composition H5N4F1, thirty-six isomers that may occur in mammals plus four isomers eventually cropping up in glyco-engineered plants were bio-synthetized from scaffold glycans prepared from human IgG, bovine fibrin and beans by the use of recombinant glycosyl-transferases (Figure S1). The choice of this type of fucosylated complex type glycan was guided by reports on the role of fucosylation for synapsin expression ^34^ together with observation that brain N-glycans appear with structures other than the standard biantennary core fucosylated oligosaccharides found on IgG ^23, 28, 35, 36^.

The many isomeric structures (Figure 1) need to be baptized and we hope to find the reader inclined to accept the herein applied system, which allows to fully define structures with short text strings naming the terminal residues as explained in ^31^ and in the Supplementary information that also provides a JAVA applet for translation into IUPAC code.

### Preparation of standards I – individual structures

The bio-synthesis of biantennary glycans with either core fucose, Lewis X (LeX) or Lewis A (LeA) determinants or with the blood group H (bgH) α1,2-fucose started from isolated GnGnF^6^, A^4^GnF^6^, GnA^4^F^6^ and A^4^A^4^F^6^ peaks from human IgG ^37^. One route entailed immediate β1,3-galactosylation yielding the core-fucose series. De-fucosylation of these standards resulted in biantennary glycans that were individually treated with either FucT-III, which is primarily an α1,4-fucosyltransferase, FucT-IV, which forms LeX epitopes, or FucT-II, which leads to bgH type glycans (Figure S1). Another fate of the core-fucosylated glycans was to serve as scaffolds for the synthesis of α-galactosylated biantennary structures (Figure 1). A large set of hybrid-type structures was synthetized from Man5 via Man5Gn. A complex variety of reactions led to hybrid-type glycans with core or outer arm fucoses with surprising results as detailed in the following chapter. The isomer assortment in this proof-of-concept work, though large, still may be expanded, *e.g.* by glycans with blood group A and B features.

The products, when subjected to PGC-LC-ESI-MS, demonstrated the impressive selectivity of the PGC stationary phase with retention times spanning a near to 30 min window (Figure S2). Many of the isomers, in particular the blood-group H variants, eluted in close proximity and this proved problematic in case of a method with notoriously unstable retention times such as PGC-LC. We therefore aimed at simultaneous acquisition of elution times of groups of peaks on the one hand and at normalization of retention times on the other.

### Preparation of standards II – differentially isotope-labelled isomer ensembles

With elution time differences often in the sub-minute and fluctuations in the minute range, consecutive injection of standards could neither provide a concise retention time library nor a reliable comparison with actual samples. Therefore, stable isotope labelling was considered for a series reference glycans on the one and for internal standards on the other hand. Deuterium introduction at the reducing end, galactosylation with one or two residues of ^13^C_6_-galactose or use of ^13^C_2_-N-acetylated compounds allowed mass increases of 1, 6 and 8 Da and in combination a panel of well over ten different increments. So, a careful choice of the preparation scheme could equip many of the standards with individual mass labels, thus allowing unambiguous identification of otherwise isobaric compounds in a chromatographic analysis. With the intention of running several standards at once, we bio-synthetized sets of isomers with inherently different mass increments as depicted in (Figure 2, Figure S3). The key point was the use of a mixture of A^4^A^4^, A^4^A^3^, A^3^A^4^ and A^3^A^3^ – derived from human IgG as described above - with ^13^C_6_ galactose as the 3-linked terminal hexose, thereby introducing three mass levels for four (or eight in case of the bgH series) isomers (Figures 2 and 3). These were combined with NaBH_4_ reduction (Lewis A series; Figure S4), NaBD_4_ reduction (both Lewis X and Core series as they elute far apart) and ^13^C_2_ acetyl groups (bgH series) (Figure 3). Isobaric core fucose, Lewis X or Lewis A structures could be assigned via the individual standards (see above), the marked difference in abundance of 6-and 3-arm galactose in human IgG, and their distinctive MS/MS spectra (Figures S5-S7), which also impressively demonstrate fucose migration. In case of the bgH-series, the bias of the α1,2-FucT for type I chains, together with single standards likewise allowed unambiguous assignment of all peaks (Figure 3). The pronounced tendency of the 3-arm to generate a B-ion fragment in positive mode MS/MS gave additional credibility to the assignments (Figure S8).

**Fig. 2.**
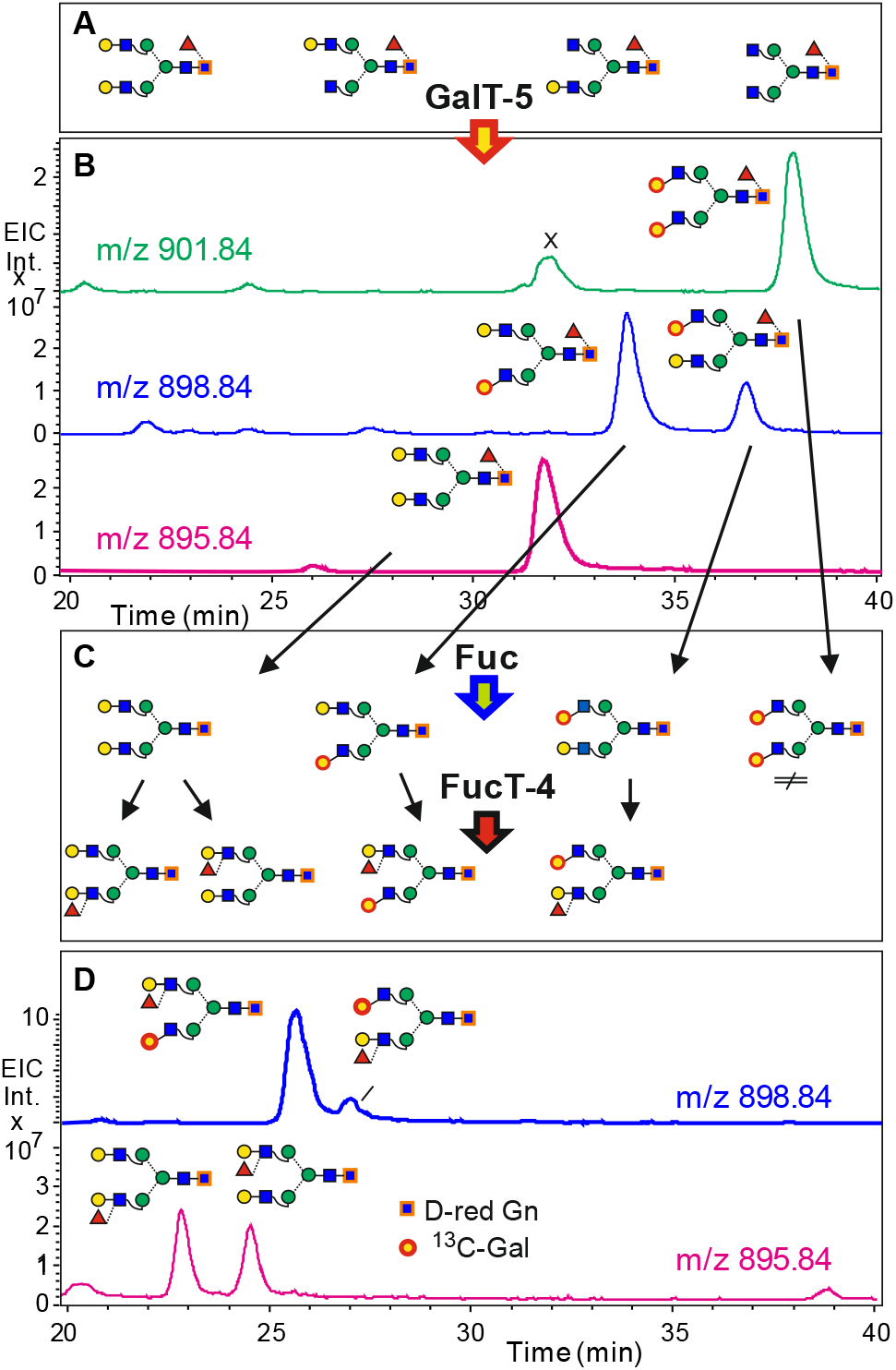
Simultaneous determination of retention times by isomer specific isotope labelling. Neutral human IgG N-glycans (**A**) were deuterium-reduced and β1,3-galactosylated by bGalT5 with UDP-^13^C_6_-galactose. The products were analyzed by PGC-LC-ESI-MS (**B**) and treated with fucosidase and FucT-4 (**C**) to generate the four LeX isomers (**D**).

**Figure 3.**
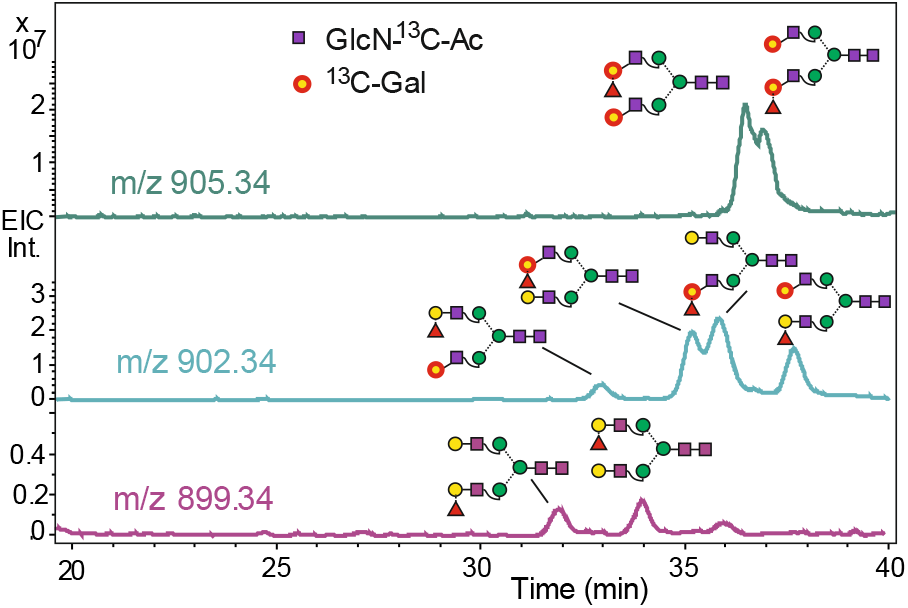
Simultaneous determination of retention times of blood group H isomers. The reaction scheme shown in Fig. 3 was conducted with N-glycans harboring four ^13^C_2_-acetyl groups and FucT-2, which acts on type 1 and type 2 chains with a roughly five-fold preference for type 1 acceptor chains.

Choice of heavy atoms for hybrid-type glycans was restricted. Man5GnF^6^bi could be equipped with a 1 Da increment only; other glycans were furnished with one ^13^C_6_-galactose residue. Having deciphered the structure of the second brain H5N4F1 peak as Man5GnF^6^bi, this glycans served as starting point for the synthesis of the isotope-labelled hybrid-type isomers. The structure was decore-fucosylated, reduced with NaBD_4_ and ^13^C_6_-galactose was subsequently introduced in either 1,4 or 1,3-linkage. The resulting structures were gently treated with α-mannosidase from jack-bean and finally with fucosyl-transferases (Figure S3). Attempts at generating the LeA determinant failed, which is in line with a recent report on the suppressive effect of bisecting GlcNAc^29^.

The conversions were at first done assuming that both b3GalT and b4GalT would faithfully transfer galactose to the 3-arm GlcNAc to generate M^3^F^2-3^bi, M^3^F^2-4^bi, M^3^(AF)bi, M^3^A^3^F^6^bi and M^3^A^4^F^6^bi. This naive assumption, however, had to be revised as b4GalT generated – in very different ratios – two products. Comparison of MS/MS spectra of the three products strongly indicated the late eluting b4GalT product of bearing the galactose on the bisecting GlcNAc (Figure 4). Experiments with Man5Gn or MGn had shown that the dominant cleavage occurs at the linkage of GlcNAc to the 3-arm α-mannose. Loss of only GlcNAc, as opposed to labelled Gal-GlcNAc disaccharide, argues for an unsubstituted 3-arm GlcNAc in compound c (Figure 4). This realization entails consequences for subsequent modifications by e.g. FucT-2 (Figure S9) and FucT-4 (see chapter on brain glycans).

**Figure 4.**
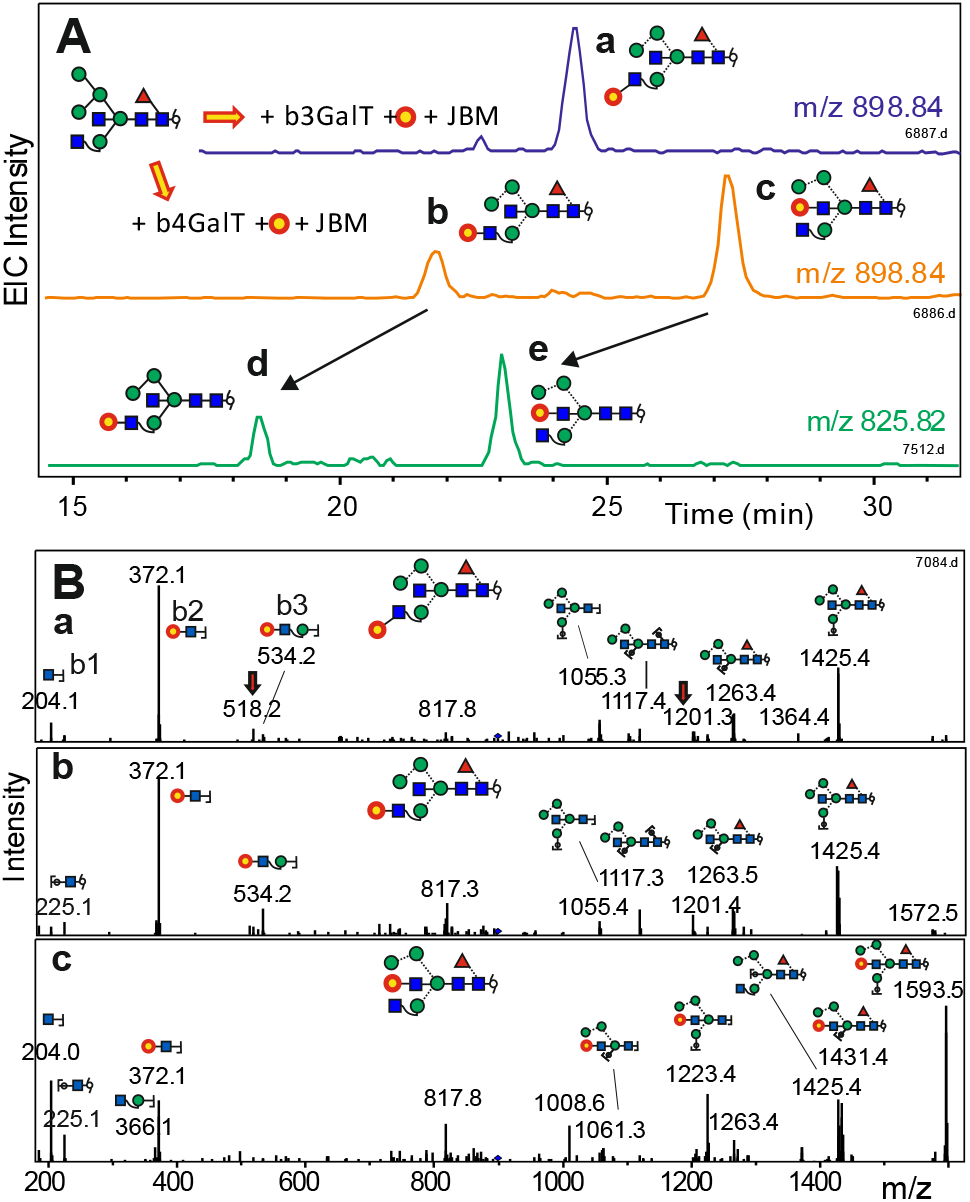
Evidence for the formation of hybrid-type N-glycans with bisecting LacNAc. Panel **A** demonstrates the effect of ^13^C_6_-galactose transfer to Man5GnF^6^bi (BD_4_-reduced) by either b3GalT or b4GalT followed by mild mannosidase digestion (compounds a, b and c) and removal of the core fucose (peaks d and e). Panel **B** shows the positive mode MS/MS spectra of peaks **a, b** and **c** (MH^+^ = 1796.68). Cartoons are only given for some y and b ions with tentative selection of possible isomers. The red arrows denote products of internal re-arrangement.

### Preparation of standards III - the time grid concept

To make a glycan retention times library useful for PGC-LC, individual runs must undergo a process akin image warping in 2D-electrophoresis, where experimental data are projected to what might be called a “mother chromatogram”. The “glucose unit” method, popular for fluorescence labelled glycans ^8, 9^ was recently adopted for PGC-LC ^38^. However, in our hands, ionization efficiency and peak shape of the isomaltose oligosaccharides of eight glucose units and above were deplorable. Besides, the likelihood of differing sorption isotherms for compounds without and with amide functions makes these simple sugars doubtful for a stationary phase as delicate as PGC. We therefore decided for a set of eight isotope labelled N-glycans with elution times covering the entire time range of the isomers in focus (Figure 5). This multi-point retention <u>**Ti**</u>me <u>**Gr**</u>id (glyco-TiGr) allowed to correct for elution time shifts, so that real elution times obtained in a particular run could be converted to virtual times (virtual minutes = *vi-min*) in an arbitrarily defined “mother chromatogram”. Interpolation of sample retention times between “glyco-TiGr” mix “sign posts” - facilitated by a dedicated Excel sheet (Supplementary Information) - generates a list of normalized retention times that can be looked up in the retention time library (Table 1). Notably, while applied to the composition H5N4F1 in this study, the very same “TiGr mix” can also be used to express elution of glycans with other masses in a manner essentially independent of the individual column and gradient conditions as demonstrated by the accidental collection of glycans with compositions other than H5N4F1 (Supplementary Table 1).

**Figure 5.**
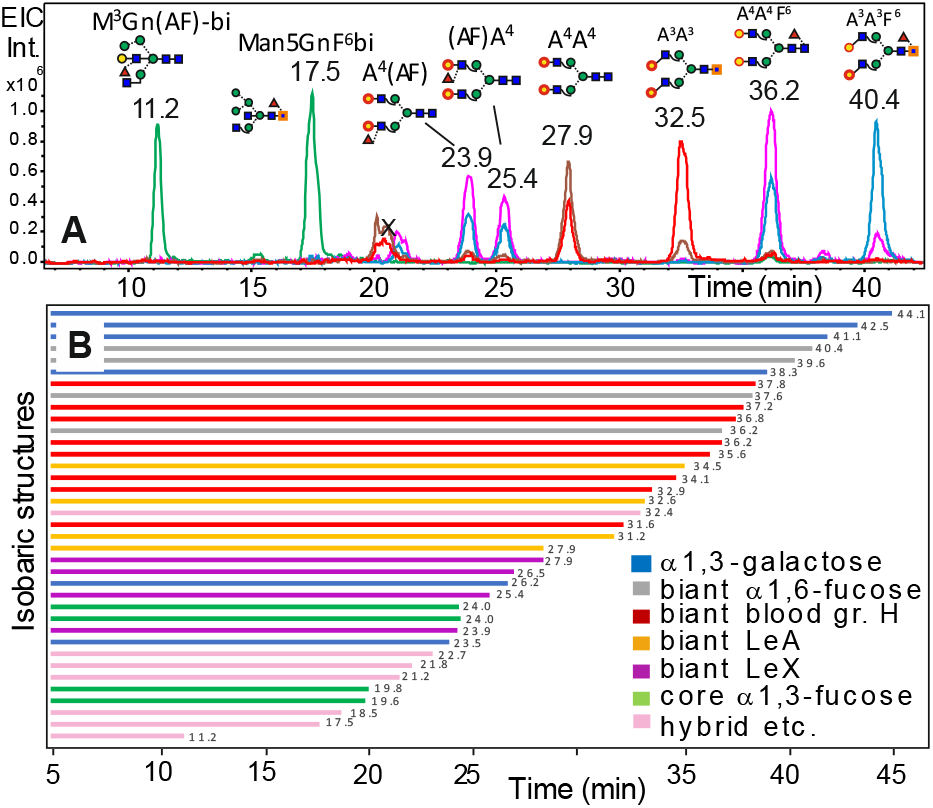
Extracted ion chromatograms for the isotope labelled internal standards defining the glyco retention time grid “glyco-TiGr” and a comprehension of the reference retention times of all glycans listed in Figure 1.

**Table 1.** Retention times of H5N4F1 isomers in the virtual “mother chromatogram”. Structure codes can be deciphered by reference to Figure 1, by use of the proglycan084 applet or reading the explanation in the Supporting Information. Two anisobaric TiGr-reference glycans are marked by “< >”. Bold print denotes structures not described so far. Retention time are given as virtual minute (vi-min).

| Proglycan code | Ret. Time (vi-min) | Proglycan code | Ret. Time (vi-min) |
| --- | --- | --- | --- |
| **M<sup>3</sup>Gn(AF)-bi** | **11.2** | A<sup>3</sup>(FA) | 31.2 |
| M<sup>3</sup>A<sup>4</sup>F<sup>6</sup>bi | 16.6 | F<sup>2-4</sup>A<sup>4</sup> | 31.6 |
| Man5GnF<sup>6</sup>bi | 17.5 | Man5An<sup>4</sup>F<sup>6</sup> | 32.4 |
| M<sup>3</sup>F<sup>2-4</sup>bi | 18.5 | < A<sup>3</sup>A<sup>3</sup> > | <32.5> |
| M<sup>3</sup>A<sup>3</sup>F<sup>6</sup>bi | 18.8 | (FA)A<sup>4</sup> | 32.5 |
| A<sup>4</sup>A<sup>4</sup>F<sup>3</sup> | 19.6 | F<sup>2-4</sup>A<sup>3</sup> | 32.9 |
| A<sup>4</sup>A<sup>3</sup>F<sup>3</sup> | 19.8 | A<sup>4</sup>F<sup>2-4</sup> | 34.1 |
| **M<sup>3</sup>GnF<sup>2-4</sup>-bi** | **21.0** | (FA)A<sup>3</sup> | 34.5 |
| M<sup>3</sup>F<sup>2-3</sup>bi | 21.2 | F<sup>2-3</sup>A<sup>4</sup> | 35.6 |
| Man5(AnF) | 21.8 | A<sup>4</sup>F<sup>2-3</sup> | 36.2 |
| **M<sup>3</sup>GnF<sup>6</sup>A-bi** | **22.7** | A<sup>4</sup>A<sup>4</sup>F<sup>6</sup> | 36.2 |
| (A<sup>3-4</sup>F<sup>3</sup>)Gn | 23.5 | F<sup>2-3</sup>A<sup>3</sup> | 36.8 |
| A<sup>4</sup>(AF) | 23.9 | A<sup>3</sup>F<sup>2-3</sup> | 37.2 |
| A<sup>3</sup>A<sup>4</sup>F<sup>3</sup> | 24.0 | A<sup>4</sup>A<sup>3</sup>F<sup>6</sup> | 37.6 |
| A<sup>3</sup>A<sup>3</sup>F<sup>3</sup> | 24.0 | A<sup>3</sup>F<sup>2-4</sup> | 37.8 |
| (AF)A<sup>4</sup> | 25.4 | A<sup>3-4</sup>GnF<sup>6</sup> | 38.3 |
| Gn(A<sup>3-4</sup>F<sup>3</sup>) | 26.2 | A<sup>3</sup>A<sup>4</sup>F<sup>6</sup> | 39.6 |
| (AF)A<sup>3</sup> | 26.5 | A<sup>3</sup>A<sup>3</sup>F<sup>6</sup> | 40.4 |
| < A<sup>4</sup>A<sup>4</sup> > | <27.9> | GnA<sup>3-4</sup>F<sup>6</sup> | 41.1 |
| A<sup>3</sup>(AF) | 27.9 | A<sup>3-3</sup>GnF<sup>6</sup> | 42.5 |
| A<sup>4</sup>(FA) | 27.9 | GnA<sup>3-3</sup>F<sup>6</sup> | 44.1 |

The criteria for choosing the TiGr mix structures were on the one hand their distribution over the elution range of the compounds of interest and on the other hand their accessibility. With regard to this economic criterion, the very late eluting standard GnA^3-3^F^6^ was not included. Certainly will further TiGr standards be added to cover the elution regions of larger and of sialylated N-glycans ^22^ (glycostore.org/showPgc).

Admittedly, not all of the fourty isomers are satisfactorily separated. But some notoriously difficult and usually ignored question can be answered at first sight. While the type of outer arm fucosylation is well accessible to negative mode MS/MS ^39^, arm location of Lewis determinants and the linkage of galactose essentially evade MS/MS strategies but are mostly unambiguously identified by retention time as seen at the example of IgA N-glycans (see below).

### The pesky Man4 issue

For Man4Gn and similar glycans, the exact structure is often not indicated. We felt uncomfortable with this anachronistic uncertainty. To resolve the issue, we conducted a limited digest of Man5Gn with jack bean α-mannosidase, which yielded two peaks. Further treatment with an α1,6-specific mannosidase eliminated the small, later eluting peak (Figure 10). The comparably fast removal of the 6-linked mannose results in a pronounced forward shift of the glycan. Thus, with the elution time of a hybrid-type Man5 glycan known, the exact arrangment of mannoses in the 6-branch of the corresponding Man4 structure can be described.

### Application to multimeric immunoglobulin A (IgA)

IgA, in particular the secretory chain present in secreted IgA, carries Lewis type glycans in addition to the core-fucosylated heavy chain structures ^19^. PGC-ESI-LC shows one large peak for the composition H5N4F1, which incidentally coelutes with the reference glycan A^4^(AF) (Figure 6). Thus, by elution time only, the outer arm fucose is assigned as Lewis X structure on the 3-arm. With retention times of essentially all other alternatives known, this can be seen as a nearly unambiguous assignment for which MS/MS only constitutes a final corroboration. Two other relatively large peaks are identified as a) the ubiquitous A^4^A^4^F^6^ and b) as (AF)A^3^, which has a type I chain and thus is not the arm isomer of the major structure. Interestingly, both the Lewis X and the core-fucose structure obviously also occur to some extent with β1,3-linked galactose, which is a structural variant difficult to seize by MS/MS and hardly ever reported for human glycoproteins. Finally, a small early eluting peak can be seen as a hybrid-type structure.

**Figure 6.**
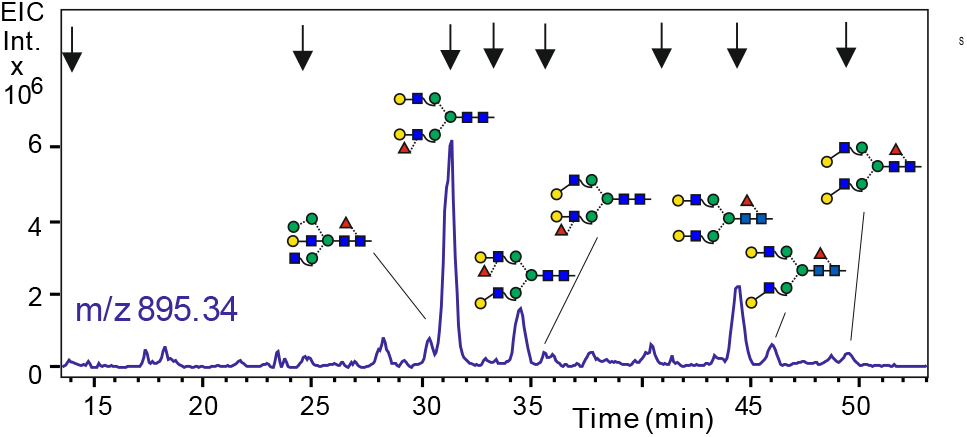
EIC for the H5N4F1 level of multimeric human IgA. At least seven isomers could be identified with reference to the mother chromatogram (Supplementary Table 2).

### Application to animal brain N-glycans

Already after having finished preparation of the four series of biantennary standards (core, LeX, LeA, bgH), we set out to compare their retention times with that found in mouse brain. The cold surprise was that none of these twenty isomers co-eluted with the three main peaks from brain. One speculation involved the presence of α1,3-galactose, a feature that was not indicated by the mass profile of mouse brain (Figure S11) and - more severe – that would cause an increase of retention ^30^ (Table 1). A further possibility allowing a H5N4F1 composition was the presence of hybrid-type glycans with bisecting GlcNAc. Indeed, the three early eluting peaks (Figure 7) co-eluted with the bio-synthetized hybrid-type references Man5GnF^6^bi, the galactosylated Man4 version of the latter and a version with LeX fucose. Negative mode ESI spectra confirmed the LeX fucose for peak 1 and core fucose for the other two peaks (Figure 12).

**Figure 7.**
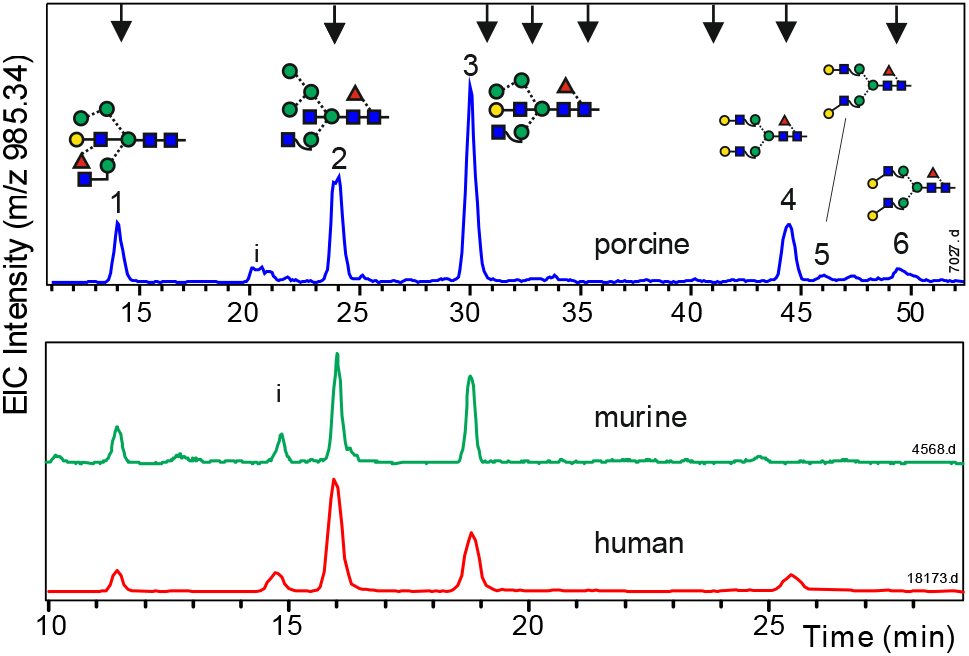
N-glycans of H5N4F1 composition from brains of various species. Arrows at the top indicate elution positions of the glyco-TiGr standard mix in the porcine brain sample. “i” denotes a peak resulting from in-source decay of a difucosylated glycan. Notably, peak 3 contains a galactosylated bisecting GlcNAc (bisecting LacNAc) and peak 1 a bisecting Lewis X determinant. Peaks 4 to 6 are supposed to result from insufficient bleeding.

This could have been the end of the story, if we would not have been sensitized by the galactose incorporation experiment with bGalT4 that gave two products (Figure 4). In fact, brain peak 3 perfectly coe-luted with product **c** and was insensitive to galactosidase, which was already observed in a previous work on bisecting LacNAc ^40, 41^. Likewise, the structure exhibited the informative preferential loss of one terminal GlcNAc in positive mode MS/MS (Figures S13 and S14). Thus, peak 3 was identified as bearing a bisecting LacNAc. The isomer with a substituted 3-arm was not observed.

The earliest eluting brain N-glycan was at first interpreted as having the structure M^3^(AF)bi, i.e. a glycan with a LeX determinant on the 3-arm GlcNAc. Its positive mode MS/MS, however, exhibited features (m/z 204 and 1586) indicative of a free 3-arm GlcNAc (Figure 8). Further proof for the presence of bisecting LeX was provided by digestion with a 3/4-specific fucosidase, which generated a peak that co-eluted i) with peak **e** in the galactose incorporation experiment (Figure 4) and ii) the peak resulting from core-de-fucosylation of brain peak 3 (Figure 8). Thus, peak 1 harbored a glycan termed M^3^Gn(AF)-bi, in which the bisecting GlcNAc was fully elaborated to a Lewis X trisaccharide, whereas the corresponding isomer M^3^(AF)bi could not be found.

**Figure 8.**
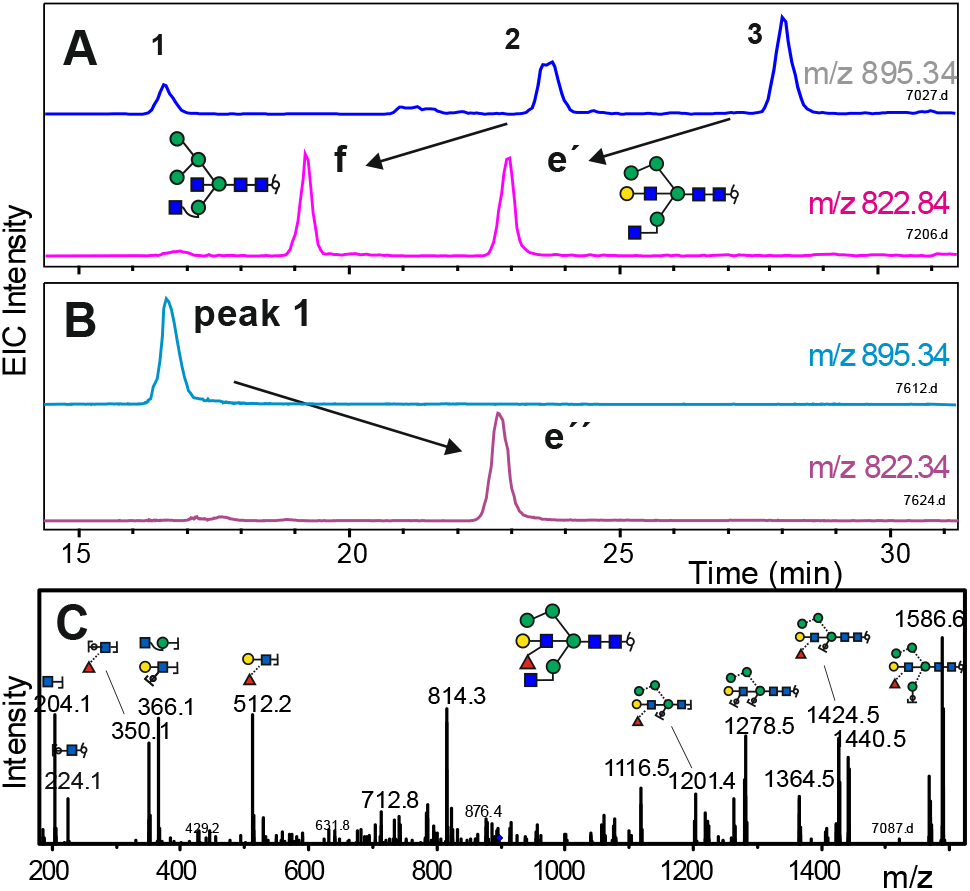
Evidence for bisecting LacNAc and bisecting Lewis in brain N-glycans. Panels **A** shows the effect of digestion of a porcine brain sample with bovine kidney fucosidase, which removes core-fucose. The disproportionate height of peak **f** arises from already present Man5Gnbi. Panel **B** depicts the effect of a1,3/4-specific fucosidase on isolated peak 1. Panel **C** shows the MS/MS spectrum of peak 1 from human brain showing the strong loss of single GlcNAc typical for free 3-arm GlcNAc.

To sum up, H5N4F1 N-glycans in brains from three species, including human, adopt three hybrid type structures of which, to the best of our knowledge, only the structure of peak 2 has been previously found in brain ^28^. The other two peaks present a bisecting LeX determinant and a bisecting LacNAc motif. These novel structures may be a foretaste of the peculiarity of brain N-glycan structures. In the absence of reference compounds, we reject the temptation for quick and all too possibly false speculations on the structures of peaks of other mass levels (Figure S15).

With the now refined view on galactose containing hybrid-type N-glycans, the structure of a minor glycan in multimeric IgA was likewise assigned as harboring a bisecting LacNAc (Figure 6). The porcine sample additionally contained diantennary N-glycans that very likely originated from contaminating blood.

### Remarks on MS/MS fragmentation

Compared to positive-mode MS/MS, negative-mode MS/MS yields more informative spectra ^23, 39^ and prevents deception by gas-phase re-arrangements ^42^. With structures already known, the positive mode fragments, revealed potential for supporting isomer identification despite a certain inclination for fucose migration between antennae and also core, as could be clearly demonstrated with asymmetrically isotope labelled glycans (Figure S5-S8). The fragment containing the GlcNAc-linked to the 3-branch mannose appeared with much higher intensity in diantennary glycans. This feature proved helpful in exposing the substitution status of the 3-arm GlcNAc of bisecting glycans (Figure 4). B-ions containing either core-fucose (Fuc-GlcNAc-ol, m/z 370) or antennary fucose (Fuc-GlcNAc, m/z 350) were prominent only in hybrid-type but not in diantennary glycans. This indicates that the yield of potential fragments is influenced by structural features rather remote from the site of fission.

LacdiNac units reveal themselves by a strong m/z 407 ion. Alfa-Gal gives rise to m/z 528.2; a m/z 674.3 ion, however, may also arise from a migrating core fucose (Figure S15). Discrimination between types of outer-arm fucosylation, however, is not provided in positive mode.

## Discussion

At the beginning of this work, the high capability of porous graphitic carbon chromatography was already known from previous studies ^17, 30, 31, 38^. The results achieved for the isobaric N-glycans containing one fucose, five hexoses and four N-acetylhexosamine residues, however, overshot our expectations. In order to secure this treasure, measures for correcting and normalizing retention times need to be developed. In contrast to the rather robust retention times in HILIC-LC, where elution times can be comfortably expressed in glucose units ^7, 19^, PGC-LC suffers from shifting elution times due to column aging (e.g. through redox/reactions with the solvent/analytes). An isomaltose oligosaccharide ladder was applied as tool to standardize retention times ^38^, but gave very poor peak shapes and insufficient coverage of the N-glycan elution range in the authors’ hands. Moreover, we consider that the lack of amide groups in oligo-glucoses could result in changes of the sorption behavior relative to N-glycans on a highly selective stationary phase such as graphitized carbon. Therefore, synthetic stable isotope-labelled N-glycans spanning the entire oligosaccharide elution range were employed as internal standards. Obviously, these glyco-TiGr standards can also be applied to N-glycans with compositions other than H5N4F1 as humbly indicated in Table S1. Likewise, the approach can be easily extended to larger, later eluting glycans simply by adding reference glycans. Population of further mass levels with a near to comprehensive collection of possible isomers would obviously result in an unparalleled isomer assignment power. Searched against a properly well-sorted virtual retention time library, the *vi-min* value of a peak at least excludes most of the possible isomers. Consultation of MS/MS spectra then merely would help to confirm an assignment or choose between the very few remaining options. The hitherto often covertly eluded distinction between different types of outer arm fucosylation becomes a child’s play with the glyco-TiGr approach. The influence of structural features on elution time on PGC has been observed, reported and exploited in recent years ^22, 23, 38^. Full use of this rich resource becomes possible only with deci-minute sharp definition of retention times together with a comprehensive list of possible isomers as introduced here for one, albeit important and very densely populated, mass level.

The focus of the current work on the H5N4F1 composition level revealed the potential of the herein introduced approach and surfaced several hitherto undescribed structures. In the IgA sample at least seven H5N4F1 isomers can be seen with an unambiguous identification of a LeX determinant on the 3-arm of a diantennary glycan. Notably, glycans with β1,3-linked galactose can be seen, which is – cautiously stated – a rare feature in human glycoproteins. In fact, the examples that are listed in the GlyConnect data base (glyconnect.expasy.org), arose from studies of murine or bovine samples. The structure (AF)A^3^ has not yet been fed into this data base.

Three prominent H5N4F1 N-glycan isomers were found in brains of mouse, pig and humans. Most remarkably, one exhibited a bisecting LacNAc unit, *i.e.* galactosylated bisecting GlcNAc, or even a bisecting LeX unit. Only b4GalT, but not b3GalT5, was able to generate this exotic feature. Bisecting LacNAc has been found as a minor component in IgG ^40, 43^. Bisecting LeX has been observed in a cell line deficient of GlcNAc tranferase II ^41^. The exact structures of brain H5N4F1 peaks 1 and 3 have - to the best of the authors’ knowledge – not been reported so far. The decoration of the bisecting GlcNAc in human brain glycans may be of particular significance because GnT-III, which produces the bisecting structures, shows altered expression levels in Alzheimer disease (AD) patients ^44, 45^ and other diseases ^46^. Therefore, interest in this element experiences a renaissance ^47^.

To conclude, these examples demonstrate the power of a well populated, ideally nearly complete, retention time library of a given composition, which moreover supports to pinpoint unexpected structures. The herein laid out concept could be the starting point for a full exploitation of the outstanding shape selectivity of porous graphitic carbon and thus for a methodology that truly appreciates the amazing and probably functionally significant isomeric diversity of N-glycans. Likewise does this study spotlight the brain N-glycome as a structurally unexplored territory, with broad implications for future glycomic studies on brain disorders such as Alzheimer disease.

## Methods

### Materials

For the preparation of reference glycans from simple scaffolds recombinant glycosyltransferases such as FucT-III, FucT-IV, FucT-VIII, GlcNAcT-I and GlcNAcT-III were expressed in the baculovirus insect cell system and purified (with exceptions) by metal chelate affinity. FucT-II was purchased from Bio-techne (Minneapolis, USA), α1,3-GalT from Chemily Glycoscience (Peachtree Corners, USA). Bovine β1,4-GalT, fungal β-galactosidase, bovine kidney fucosidase, jack bean α-mannosidase were from Sigma-Aldrich (Vienna, Austria) and α1,6-mannosidae as well as α1,3/4-specific fucosidase from New England Biolabs (Ipswich, MA, USA). UDP-^13^C_6_-galactose was prepared from ^13^C_6_-galactose (Cambridge Isotope Laboratories, Tewksbury, MA, USA) with galactokinase (Sigma-Aldrich, Vienna, Austria) and recombinant galactose 1-phosphate uridylyltransferase as described ^48^. The biantennary N-glycan A^4^A^4^ with ^13^C acetyl groups was purchased from Asparia glycomics (San Sebastian, Spain).

Human brain samples were kindly provided by Dr. Lena Hirtler (Medical University, Vienna). Mouse brain was donated by Dr. Boris Ferko (then Dep. of Biotechnology, University of Renewable Resources and Life Sciences, Vienna). Pig brain was bought at a local butcher.

### Glycan preparations

N-glycans from brains, bovine fibrin, and taka amylase were prepared by pepsin digestion, glycopeptide extraction and PNGase treatment. Where desired, glycans were chemically desialylated. Human IgG and IgA were heat denatured and deglycosylated by PNGase F. Glycans were reduced with either NaBH_4_ or NaBD_4_ and eventually fractionated by PGC-LC monitored by MALDI-TOF MS ^49^. These glycans were used as scaffolds for the different glycosidases and glycosyltransferases, eventually with purification steps by PGC-LC. Details of the preparation procedures are provided in the Supplementary Information.

### LC-ESI-MS

Analytical PGC-LC-ESI-MS/MS was performed with a capillary column (Hypercarb, 100 mm × 0.32 mm, 5 μm particle size; Thermo Scientific, Waltham, MA, USA) eluted with 10 mM ammonium bicarbonate as the aqueous solvent and acetonitrile as the organic modifier ^17, 50^. Detection was performed with an ion trap instrument (amaZon speed ETD; Bruker, Bremen, Germany) equipped with the standard ESI source linked to a Thermo Ultimate 3000 UPLC system. Samples were measured in positive mode with data-dependent acquisition. Data interpretation was done with Data Analysis 4.0 (Bruker, Bremen, Germany) and GlycoWorkBench ^51^.

## Supporting information

Supplemental Methods, Tables, Figures and an Explanation of N-glycan nomenclature

## Supporting Information

Supporting Methods, Tables and Figures (Helm_TiGr_SI.pdf) as far as technical contraints will allow: TiGr calculation (Helm_Vi-min.xls) Structure representation aid (Proglycan0814.jar)

## Acknowledgments

This work was supported by the Austrian Science Fund (Doctoral Program BioToP – Molecular Technology of Proteins (W1224), BrainProt P22274) and the European Commission (Newcotiana 760331). We thank Dr. Lena Hirtler for providing human brain samples. We owe special tribute to Wolfhart Janu (né Freinbichler) for creating the proglycan Java applet.

## Author contributions

J.H. C.G., M.P. and F.A. conceived the study. J.H., A.T. J.U., J. F., L.N. and D.S. performed enzyme expressions and glycan preparations. J.H., C.G., D.M., L.N. and M.P. conducted the analytical experiments. J.H. and F.A. evaluated the data and composed the manuscript. All authors reviewed the manuscript.

## Data availability

The data supporting the results of this study is available within the article and its Supplementary Information files. Access to MS raw data files will be provided upon request.

## Notes

The authors declare no competing financial interest.

## Notes

### Competing Interest Statement

The authors have declared no competing interest.

### Summary of Updates

Only change: clear distinction between previous affiliation and current address.

