## Supplemental Methods, Tables, Figures and an Explanation of N-glycan nomenclature for "Time grid-based isomer specific N-glycan analysis and detection of bisecting Lewis X in human brain"

Content:

I. Supplementary methods

II. Supplementary tables and figures

III. Exposition of the N-glycan nomenclature system "proglycan" used in this work

**Current addresses:**
AT, Institute of Science and Technology Austria, 3400 Klosterneuburg. JU, BIOMAY AG, 1190 Wien Austria. JF VelaLabs GmbH, 1230 Wien, Austria. DM Boehringer Ingelheim, 1120 Wien, Austria. LN BakerHicks AG; 1120 Wien, Austria. MP Delft University of Technology, Delft, The Netherlands.

### Supplementary Materials and Methods

#### Materials

Mouse brains were kindly donated by Dr. Boris Ferko (now Research Institute of Animal Production (RIAP) in Nitra, SK). Porcine brain was obtained from a local butcher. Samples from two human brains were kindly provided by Dr. Lena Hirtler (Medical University, Vienna, AT).
The biantennary N-glycan $A^4A^4$ with $^{13}C_2$ acetyl groups was purchased from Asparia glycomics (San Sebastian, Spain). $^{13}C_6$-galactose was purchased from Cambridge Isotope Laboratories (Tewksbury, MA, USA).

#### N-glycan preparation from human and mouse brains

About 500 mg brain tissue (human or mouse brain) was cut into tiny pieces with the help of a scalpel. The pieces were transferred into a Falcon tube and mixed with 100 mM ammonium bicarbonate buffer containing 15 mM dithiothreitol (DTT) and 2.5 % sodium dodecyl sulfate in a total volume of 3.5 ml. After homogenization with an Ultra-Turrax T25 disperser the homogenate was boiled at 95°C for 5 min in order to eliminate enzymatic activities. After centrifugal clarification the supernatant was mixed with four volumes of methanol, one volume of chloroform and three volumes of water in the given order. After centrifugation of the mixture for 3 min at 2500 rpm, the upper phase was removed and another four volumes of methanol was added. After another centrifugation, the pellet was washed with methanol. The dried pellet was taken up in 50 mM ammonium acetate of pH 8.4 containing 1.25 M urea. For N-glycan release 2.5 U of PNGase F was added and the resulting mixture was incubated overnight at 37°C. This reaction mixture was loaded onto a Hypersep C18 cartridge (25 mg; Thermo Scientific, Vienna) that had been previously washed with 500 µl of methanol, then 500 µl 50 % acetonitrile in 65 mM ammonium formate buffer (pH 3.0) and equilibrated twice with 500 µl 0.1 % formic acid. The flow-through of the applied sample was collected and subjected to centrifugal evaporation. Reduction of the glycans was carried out in 1 % sodium borohydride at room temperature overnight. Desalting was performed using HyperSep Hypercarb solid-phase extraction cartridges (25 mg) (Thermo Scientific, Vienna).

##### N-glycan preparation from pig brain, bovine fibrin and white beans

N-glycans from pig brain and white beans (ca. 50 g) were prepared by pepsin digestion, (glyco)-peptide extraction on a cation exchange resin and PNGase A treatment as previously described [1]. Peptic glycopeptides from bovine fibrin were desialylated in 50 mM sulfuric acid for 1 h at 80 °C. After neutralization with NaOH and centrifugation the glycopeptides were enriched by gel filtration and deglycosylated [1]. Reduction and final purification were done as described above.

#### N-glycan preparation from purified glycoproteins

In the case of human IgG, 100 mg glycoprotein was dissolved in 10 mL 50 mM ammonium acetate (pH 8.4) and brought to 95 °C for 10 min. After cooling, 0.2 mU peptide:N-glycosidase F was added and digestion was carried out overnight at 37 °C. Subsequently, the released N-glycans were acidified, reduced and recovered by passage over Hypersep C18 cartridge as described above for small scale brain glycan preparations. Human colostrum IgA (1 mg) was treated similarly.

IgG N-glycans were in part fractionated by semi-preparative HPLC on a Hypercarb column as described previously [1]. The major components $GnGnF^6$, $A^4GnF^6$, $GnA^4F^6$ and $A^4A^4F^6$ eluted in this order [2] and could be obtained as isolated fractions for the preparation of defined isomers.

#### Glycosyltransferases and glycosidases

Recombinant human α1,2-fucosyltransferase (FucT-II) was purchased from Bio-techne (Minneapolis, USA), recombinant bovine α1,3-galactosyltransferase from Chemily Glycoscience (Peachtree Corners, USA) and bovine milk β1,4-galactosyltransferase from Sigma-Aldrich (Vienna, Austria). Bovine kidney fucosidase from bovine kidney and jack bean mannosidase were purchased from Sigma-Aldrich (Vienna, Austria). The β1,4-specific galactosidase from *Aspergillus oryzae* was prepared as described [3]. α1,3/4-specific fucosidase and α1,6-specific mannosidase were purchased from NEB (Ipswich, MA, USA).

Human fucosyltransferase III (FucT-III) lacking the N-terminal 34 amino acids, human fucosyltransferase IV (FucT-IV) lacking the N-terminal 172 amino acids, fucosyltransferase VIII (FucT-VIII) from *Bos taurus* lacking the N-terminal 30 amino acids, human β1,3-galactosyltransferase (b3GalT5) lacking the N-terminal 29 amino acids, *N*-acetylglucosaminyltransferase I (GnT-I) from rabbit lacking the N-terminal 105 amino acids and human *N*-acetylglucosaminyltransferase III (GnT-III) lacking the N-terminal 22 amino acids were recombinantly expressed with a N-terminal $His_6$-tag using a pVT-Bac vector and the baculovirus insect cell system as described previously [4] . The enzymes were purified by metal chelate chromatography. The buffer was changed to 25 mM Tris/HCl pH 7.4 supplemented with 100 mM NaCl and the volume was reduced to less than 1 mL with an Amicon 10 kDa cut-off membrane (SigmaAldrich, Vienna). The enzymes were directly used after purification or stored at 4 °C.

#### Preparation of $^{13}C_6$-UDP-Galactose

UDP-$^{13}C_6$-galactose was prepared by incubation of $^{13}C_6$-galactose with galactokinase (Sigma-Aldrich, Vienna, Austria). Galactose-1-phosphate was converted to the nucleotide sugar in the presence of UDP-glucose by human galactose-1-phosphate uridylyltransferase, which was recombinantly expressed in *Escherichia coli* BL21 and purified via its His6-tag. The UDP-$^{13}C_6$-galactose was finally purified by PGC chromatography with a slightly alkaline buffer as described [5].

#### Biosynthesis of glycan structures

The reduced and dried glycans, prepared as described above, were used as scaffolds for the different glycosidases and glycosyltransferases. The non-labelled standards were biosynthesized according to Supplementary Fig. 1 and the isotope-labelled glycan standards were biosynthesized according to Supplementary Fig. 3. Fucosyltransferases were used with 1 mM GDP-fucose and 10 mM MnCl2 in a total volume of 50 µL in 25 mM Tris/HCl pH 7.4 + 100 mM NaCl. Bovine β-1,4-galactosyltransferase and human β-1,3-galactosyltransferase were used with 1 mM UDP-galactose or 13C6-UDP-galactose (or 1 mM UDP-GalNAc) and 20 mM MnCl2 for bovine β-1,4-galactosyltransferase and 2 mM MnCl2 for human β-1,3-galactosyltransferase in a total volume of 50 µL in 25 mM Tris/HCl pH 7.4 + 100 mM NaCl. N-acetylglucosaminyltransferases 1 and 3 were used with 1 mM UDP-GlcNAc and 20 mM MnCl2 in 50 mM Tris/HCl pH 7.4. α1,3-galactosyltransferase was used with 1 mM UDP-galactose or 13C6-UDP-galactose and 10 mM MnCl2 in 50 µL of 25 mM Tris/HCl pH 7.4 + 100 mM NaCl. β-galactosidase from *Aspergillus oryzae* and fucosidase from bovine kidney were used in 0.1 M phosphate citrate buffer pH 5.0, α-mannosidase from jack bean was used in sodium acetate buffer pH 4.5 supplemented with 5 mM $ZnCl_2$. α1,3/4-specific fucosidase and α1,6-mannosidase were used according to the manufacturer´s protocol.

All enzyme reactions were carried out overnight at 37 °C. After each enzyme-reaction step, the reaction mixtures were purified using carbon solid phase cartridges (Multi-Sep Hypercarb 25 mg, Thermo Scientific, Vienna) and the completeness of the enzymatic reaction was checked using PGC-

LC-ESI-MS before further enzymatic processing. After the last enzymatic processing step, the finished glycan standards were again purified using PGC-SPEs, dried by centrifugal evaporation, taken up in a small volume of HQ-water and stored at -20°C until measurement.

#### Mass spectrometric methods

The purified samples were loaded on a PGC-column (100 mm × 0.32 mm, 5 µm particle size, Thermo Scientific, Waltham, MA, USA) with 10 mM ammonium bicarbonate as the aqueous solvent A and 80 % acetonitrile in solvent A as solvent B. The gradient was as followed: 0 – 4.5 min 1 % B, from 4.5 – 5.5 min 8 % B, from 5.5 – 58 min 8 - 22 % B, from 58 – 62.5 min 22 - 30 % B, from 62.5 – 63.5 min 68 % B, followed by an equilibration period at 1 % B from 64 – 70 min. The flowrate was 6 µL/min. Detection was performed with an ion trap mass spectrometer equipped with the standard ESI source directly linked to the Thermo Ultimate 3000 UPLC system. MS-scans were recorded from 400 – 1600 m/z. All samples were measured with an ion trap instrument (amaZon speed ETD; Bruker, Bremen, Germany) in positive mode or, in the case for some brain glycan runs, in negative mode. Standard source settings (capillary voltage 4.5 kV, nebulizer gas pressure 0.5 bar, drying gas 5 L/min, 200 °C) were used. Instrument tuning was optimized for a low mass range (around 1500-2000 Da molecules). MS/MS was carried out in data-dependent acquisition mode (switching to MS/MS mode for eluted peaks). Data interpretation was done with DataAnalysis 4.0 (Bruker, Bremen, Germany).

#### Supplementary Table:

**Supplementary Table 1** Vi-min values of N-glycans of various compositions. This collection of isomers does by no means claim to be comprehensive.

| **H5N4** Proglycan-code | vi-min |
|---|---|
| M$^3$A$^4$bi | 11.9 |
| Man5Gnbi | 13.0 |
| M$^3$A$^3$bi | 13.7 |
| M$^3$GnA-bi | 15.7 |
| Man5An | 23.4 |
| A$^4$A$^4$ | 27.9 |
| A$^4$A$^3$ | 29.3 |
| A$^{3\text{-}4}$Gn | 30.4 |
| A$^3$A$^4$ | 31.8 |
| A$^3$A$^3$ | 32.5 |
| GnA$^{3\text{-}4}$ | 33.4 |
| A$^{3\text{-}3}$Gn | 35.1 |
| GnA$^{3\text{-}3}$ | 35.9 |

| **H5N4F2** | |
|---|---|
| (AF)(AF) | 20.9 |
| (FA)(FA) | 32.4 |
| (F$^{2\text{-}3}$)(F$^{2\text{-}3}$) | 43.4 |
| (F$^{2\text{-}4}$)(F$^{2\text{-}4}$) | 43.6 |

| **H5N4F1** Proglycan-code | vi-min | **H5N4F1** Proglycan-code | vi-min |
|---|---|---|---|
| M$^3$Gn(AF)-bi | 11.2 | A$^3$(FA) | 31.2 |
| M$^3$A$^4$F$^6$bi | 16.6 | F$^{2\text{-}4}$A$^4$ | 31.6 |
| Man5GnF$^6$bi | 17.5 | Man5An$^4$F$^6$ | 32.4 |
| M$^3$(F$^{2\text{-}4}$)bi | 18.5 | (FA)A$^4$ | 32.5 |
| M$^3$A$^3$F$^6$bi | 18.8 | F$^{2\text{-}4}$A$^3$ | 32.9 |
| A$^4$A$^4$F$^3$ | 19.6 | A$^4$F$^{2\text{-}4}$ | 34.1 |
| A$^4$A$^3$F$^3$ | 19.8 | (FA)A$^3$ | 34.5 |
| M$^3$Gn(F$^{2\text{-}4}$)bi | 21.1 | F$^{2\text{-}3}$A$^4$ | 35.6 |
| M$^3$(F$^{2\text{-}3}$)bi | 21.2 | A$^4$F$^{2\text{-}3}$ | 36.2 |
| Man5(An$^4$F$^3$) | 21.8 | A$^4$A$^4$F$^6$ | 36.2 |
| M$^3$GnF$^6$A-bi | 22.7 | F$^{2\text{-}3}$A$^3$ | 35.6 |
| (A$^{3\text{-}4}$F$^3$)Gn | 23.5 | A$^3$F$^{2\text{-}3}$ | 37.2 |
| A$^4$(AF) | 23.9 | A$^4$A$^3$F$^6$ | 37.6 |
| A$^3$A$^4$F$^3$ | 24 | A$^3$F$^{2\text{-}4}$ | 37.8 |
| A$^3$A$^3$F$^3$ | 24 | A$^{3\text{-}4}$GnF$^6$ | 38.3 |
| (AF)A$^4$ | 25.4 | A$^3$A$^4$F$^6$ | 39.6 |
| Gn(A$^{3\text{-}4}$F$^3$) | 26.2 | A$^3$A$^3$F$^6$ | 40.4 |
| (AF)A$^3$ | 26.5 | GnA$^{3\text{-}4}$F$^6$ | 41.1 |
| A$^3$(AF) | 27.9 | A$^{3\text{-}3}$GnF$^6$ | 42.5 |
| A$^4$(FA) | 27.9 | GnA$^{3\text{-}3}$F$^6$ | 44.1 |

**Supplementary Table 2. Isomer assignment of IgA N-glycans by the glyco-TiGr approach.** Experimental retention times of the peaks from Figure 6 are converted by linear interpolation using an Excel routine (Supplementary information). Values for virtual minutes (*vi-mins*) are compared to Table 1. Structures in bracket denote highly unlikely isomers with incidentally very similar retention time.

| **Peak Nr.** | **Exp.Rt.** (min) | ***Vi-min*** | **Closest hit** | **Δ** (min) | **Structure** |
|---|---|---|---|---|---|
| 1 | 30.2 | 22. | 22.7 | 0.2 | $M^3GnF^6A$-bi |
| 2 | 31.2 | 23. | 23.9 | 0 | $A^4$(AF) |
| 3 | 34.5 | 26. | 26.5 | 0.2 | (AF)$A^3$ |
| 4 | 35.7 | 27. | 27.9 | 0 | $A^3$(AF), [($A^4$(FA)] |
| 5 | 44.3 | 36. | 36.2 | 0 | $A^4A^4F^6$,[$A^4F^{2\text{-}3}$] |
| 6 | 46.0 | 37. | 37.6 | 0 | $A^4A^3F^6$ |
| 7 | 49.4 | 40. | 40.4 | 0 | $A^3A^3F^6$ |

Peaks with two or three fucose residues were arbitrarily identified as either (AF)(AF) or as $A^4$(AF) and (AF)$A^4$. Judging from the shortened retention time compared to the doubly fucosylated structures and the occurrence of core- and Lewis X-fucose in other structures from the sample, the triply fucosylated structure at 39.0 min most likely resembled (AF)(AF)$F^6$ (Figure S6).

#### Supplementary Figures

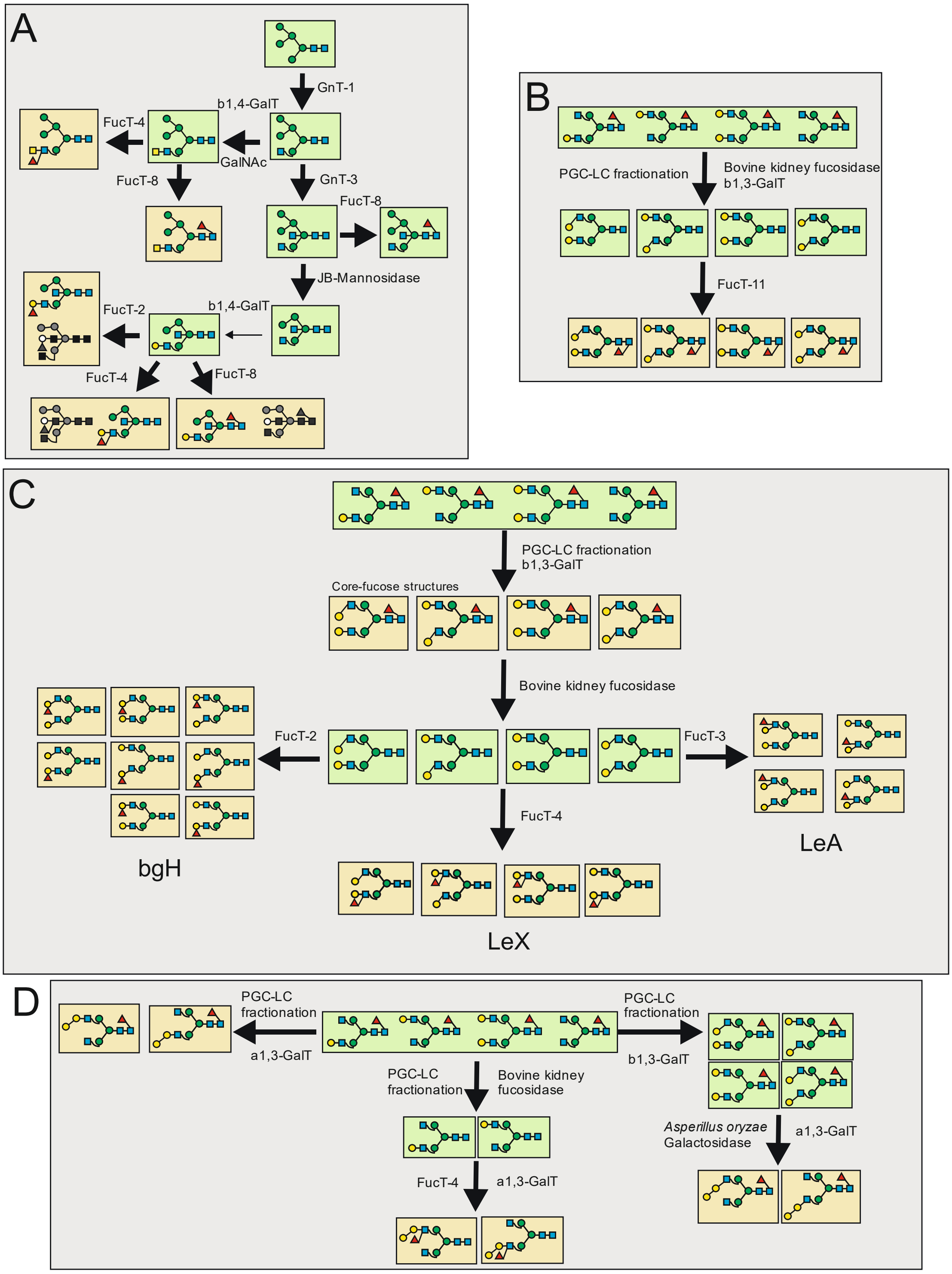

**Supplementary Fig. 1** Pathways for biosynthesis of individual N-glycan structures with the composition H5N4F1; Panel **A** describes the biosynthesis of hybrid-type structures with Man5 from beans as starting point; Structures drawn in grey scales were unanticipated results. Panel **B** shows the pathway for plant type core α1,3-core fucosylated structures from isolated IgG glycans as the starting point; Panel **C** gives the pathways for Lewis X, Lewis A, blood group H and core α1,6-fucosylated biantennary structures from isolated $A^4GnF^6$, $GnA^4F^6$, $A^4A^4F^6$ and $GnGnF^6$ glycans; Panel **D** shows the pathway for the α1,3-galactosylated biantennary structures.

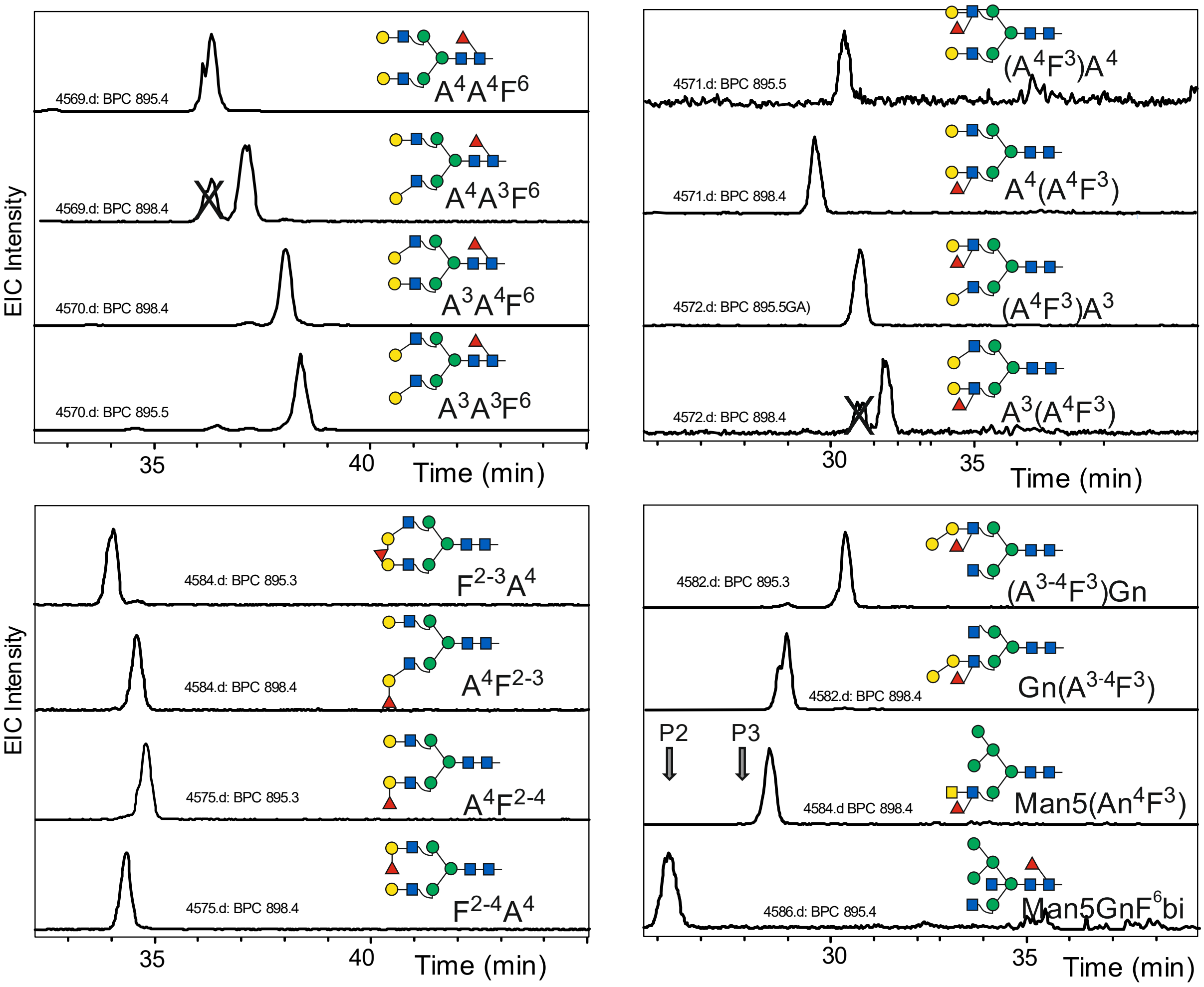

**Supplementary Fig. 2** Examples of elution positions of individual reference glycans. Performed long before acquisition of the "mother chromatogram", the elution times may deviate. In many cases, light and heavy (one $^{13}C_6$-Gal) standards were co-injected as indicated by after the run number. Grey arrows in the lower right corner denote the positions of brain H5N4F1 peaks 2 and 3.

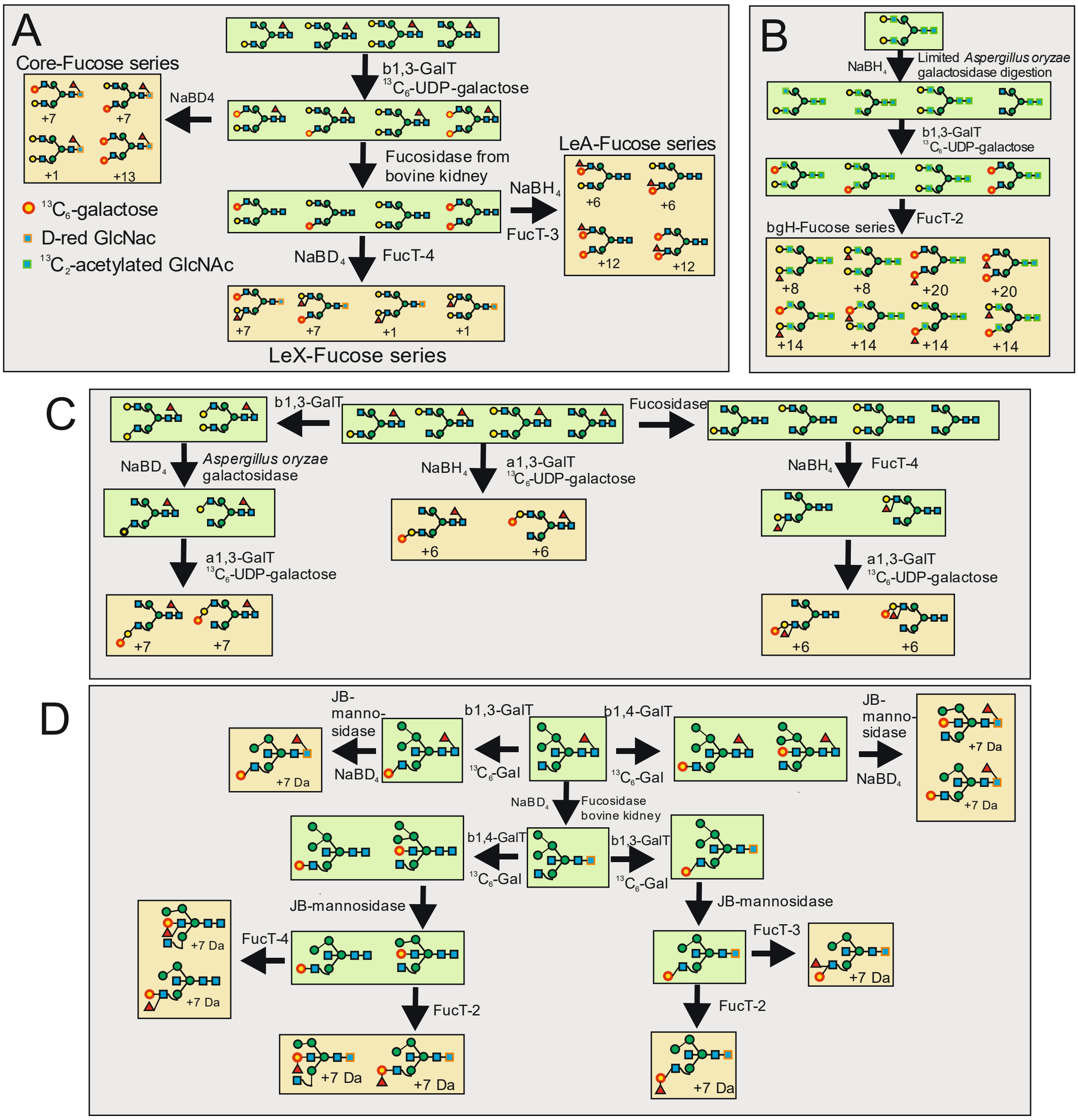

**Supplementary Fig. 3** Biosynthesis pathways for isotope labelled N-glycans with the composition H5N4F1; Panel A shows the pathways for the biosynthesis of complex-type biantennary glycans (LeX-, LeA- and core-fucose) with A4GnF$^6$, GnA$^4$F$^6$, A$^4$A$^4$F$^6$ and GnGnF$^6$ isolated from human IgG as the starting point; Panel B shows the pathway for the biosynthesis of the complex-type biantennary bgH glycans with $^{13}$C$_2$-acetylated A$^4$A$^4$ as the starting point; Panel C shows the pathways for the biosynthesis of alfa-galactosylated complex-type biantennary structures with A$^4$GnF$^6$, GnA$^4$F$^6$, A$^4$A$^4$F$^6$ and GnGnF$^6$ isolated from human IgG as the starting point; Panel D shows the pathways for the biosynthesis of hybrid-type glycans with Man5GnF$^6$bi isolated from pig brain as the starting point. Generation of a LeA determinant on hybrid-type glycans, however, failed, which reflects the recently described suppressive effect of bisecting GlcNAc on terminal modifications [6].

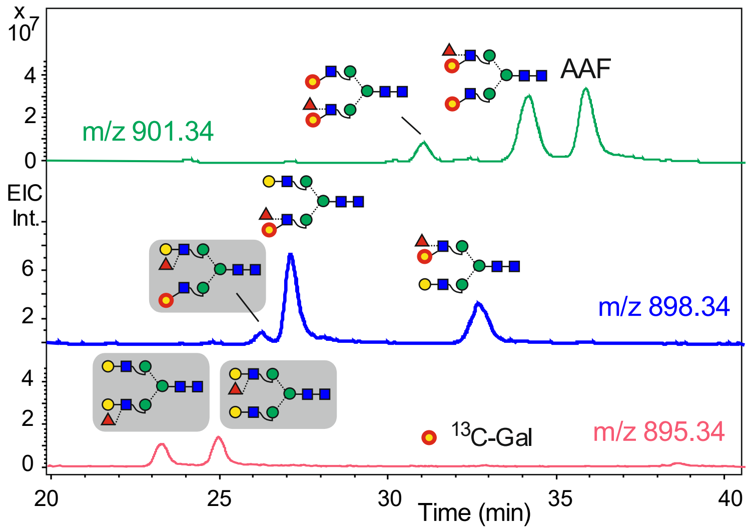

**Supplementary Fig. 4** Simultaneous determination of retention times of LeA isomers. Reaction of a $NaBH_4$ reduced, defucosylated mixture as shown in Fig. 2C with FucT-3 generated four LeA isomers. Due to the dual activity of FucT-3, LeX isomers were also formed. These alternative products are shaded in grey. The peak designated AAF arises from the overlap with the isotopic pattern of the TiGr standard $A^4A^4F^6$ (see Fig. 5).

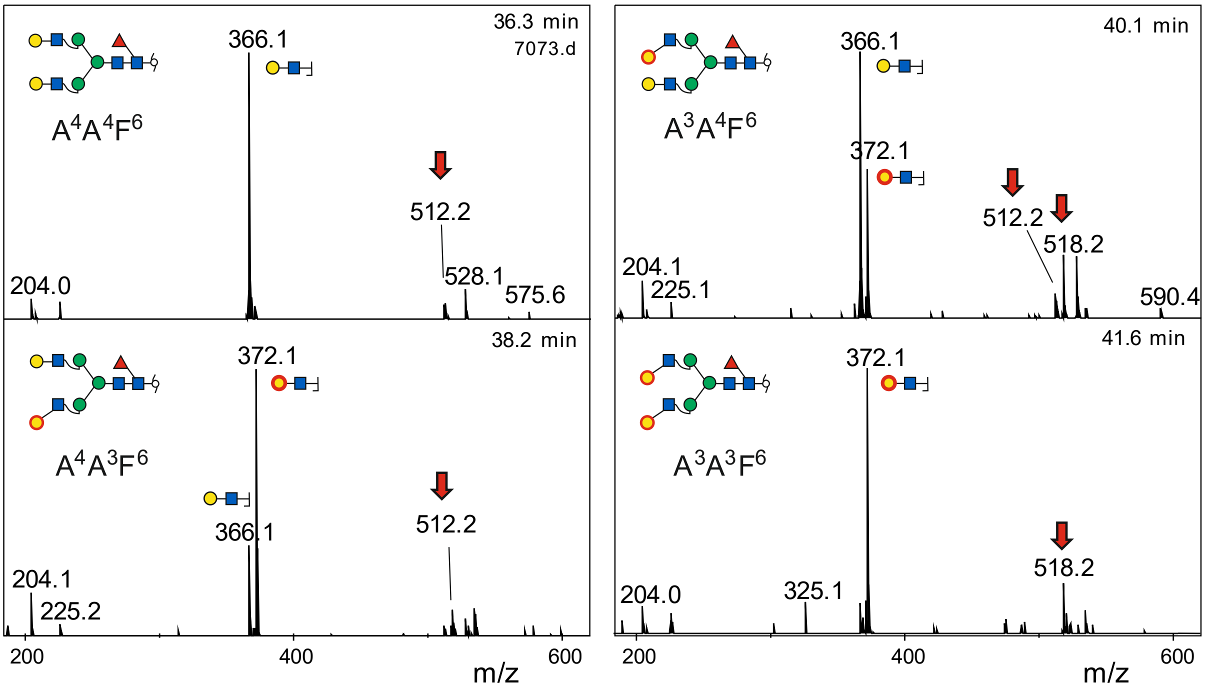

**Supplementary Fig. 5** Relevant sections of the positive mode MS/MS spectra of core fucosylated $^{13}C_6$-galactose labelled standard N-glycans. The respective EICs are shown in main text Figure 2. Note the different height of the disaccharide ions depending on their origin. Red arrows label fragments unquestionably resulting from re-arrangement.

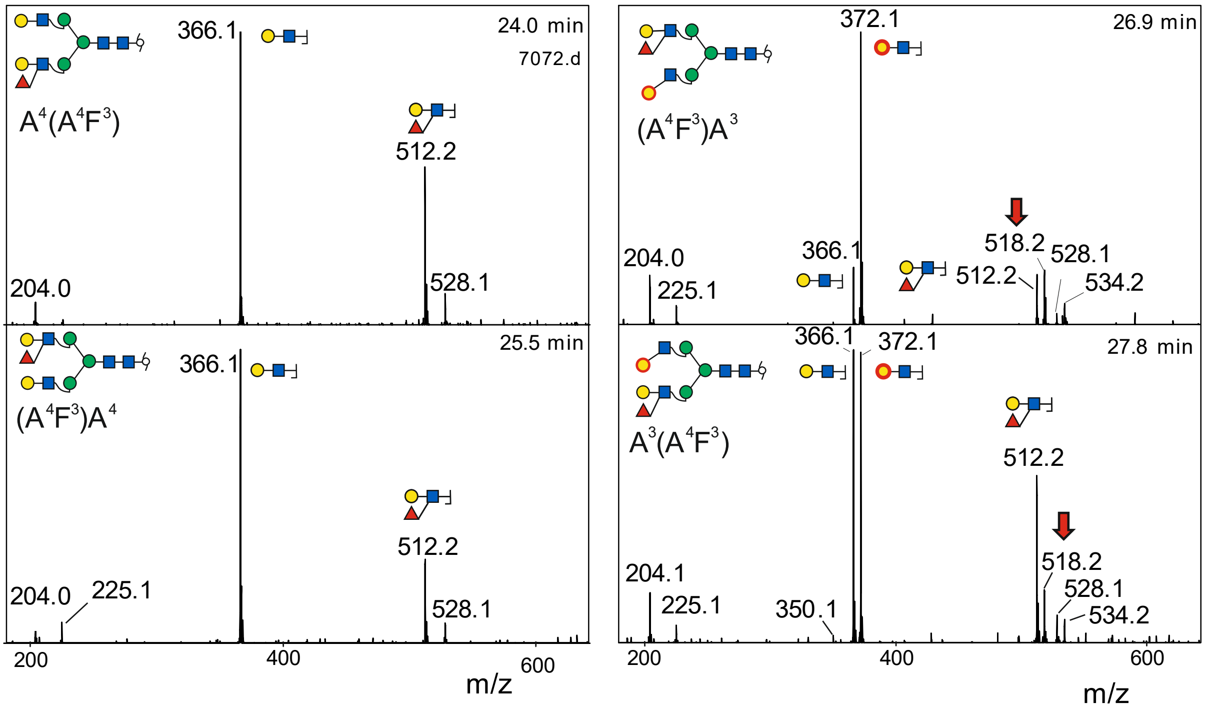

**Supplementary Fig. 6** Relevant sections of the positive mode MS/MS spectra of $^{13}C_6$-galactose labelled N-glycans with LeX determinants. The respective EICs are shown in main text Figure 2. Note the different height of the disaccharide ions depending on their origin. Red arrows label fragments unquestionably resulting from re-arrangment.

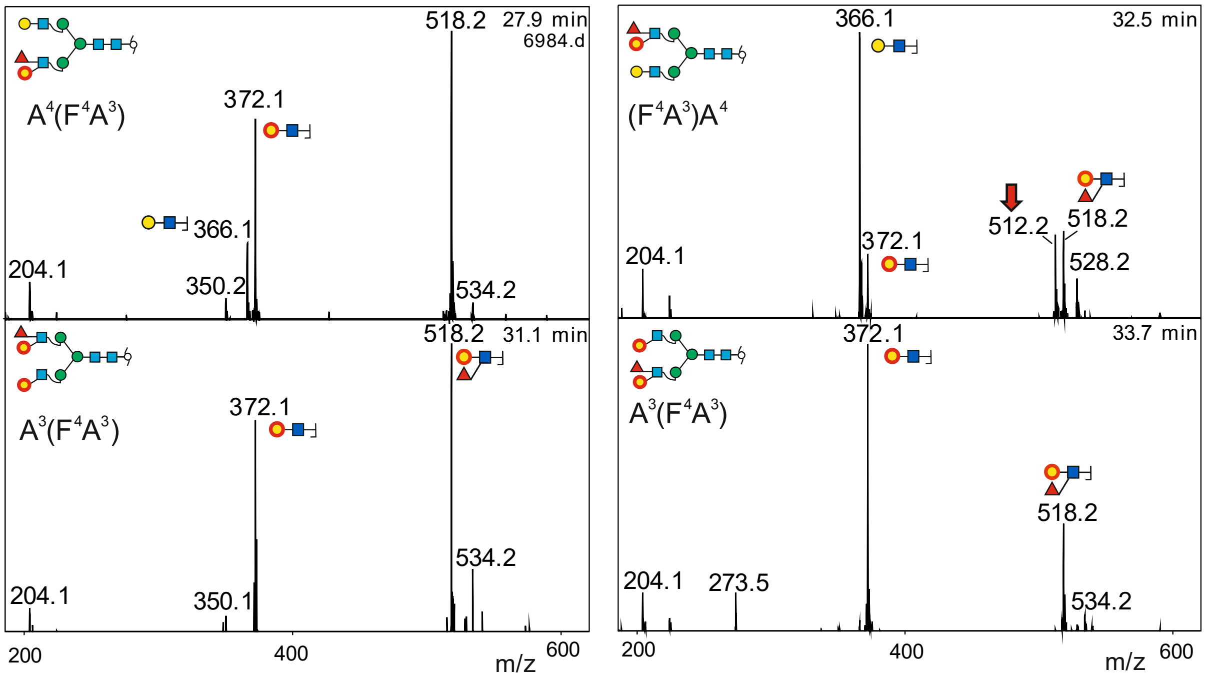

**Supplementary Fig. 7** Relevant sections of the positive mode MS/MS spectra of $^{13}C_6$-galactose labelled N-glycans with LeA determinants. The respective EICs are shown in **Supplementary Fig. 4**. Note the different height of the disaccharide ions depending on their origin. Red arrows label fragments unquestionably resulting from re-arrangement.

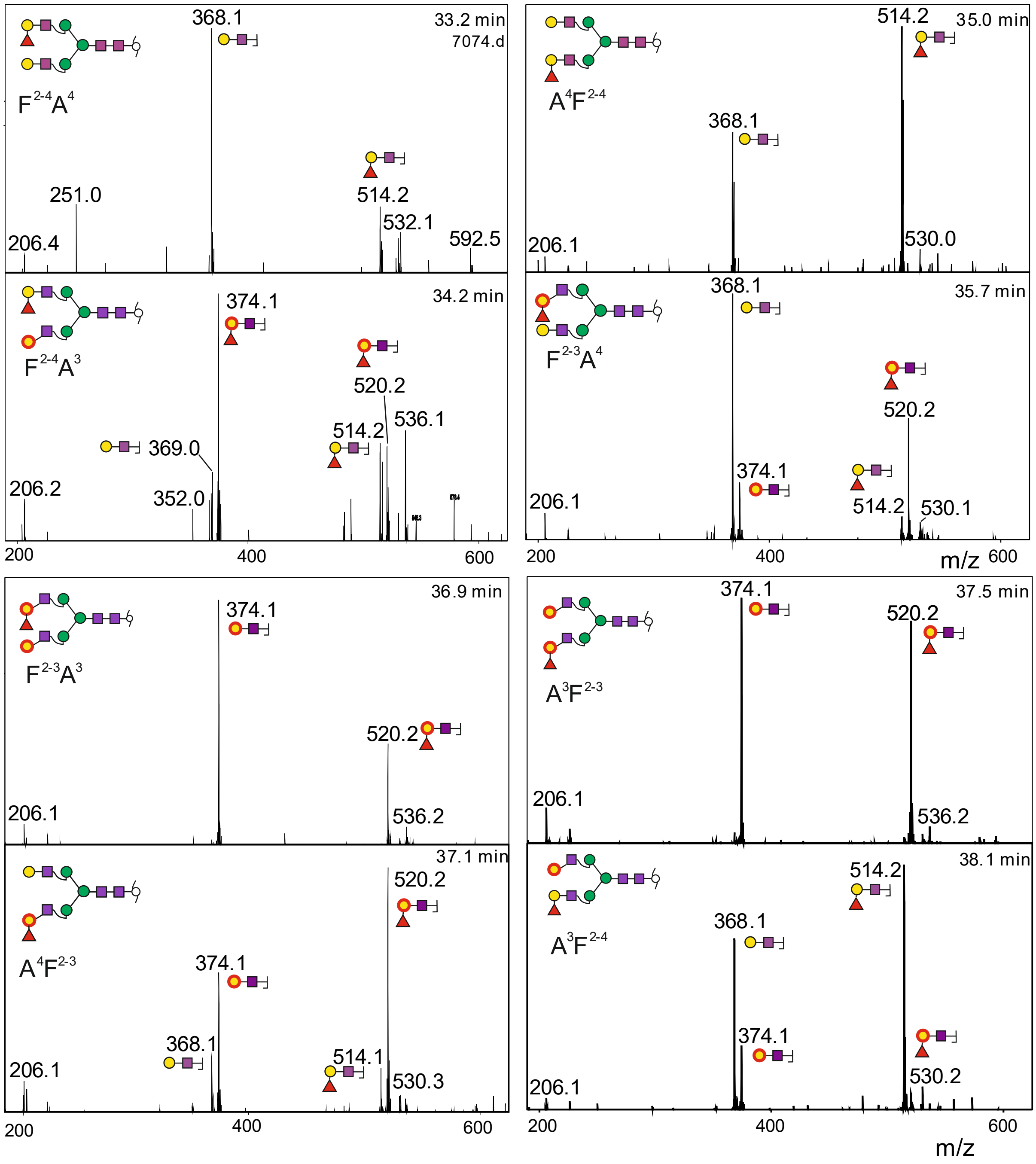

**Supplementary Fig. 8** Relevant sections of the positive mode MS/MS spectra of $^{13}C_6$-galactose (and $^{13}C_2$-acetyl) labelled N-glycans with blood group H determinants. The respective EICs are shown in main text Fig. 3. The purple squares GlcNAc with $^{13}C_2$-labelled acetyl groups. Note the different height of the disaccharide ions depending on their origin.

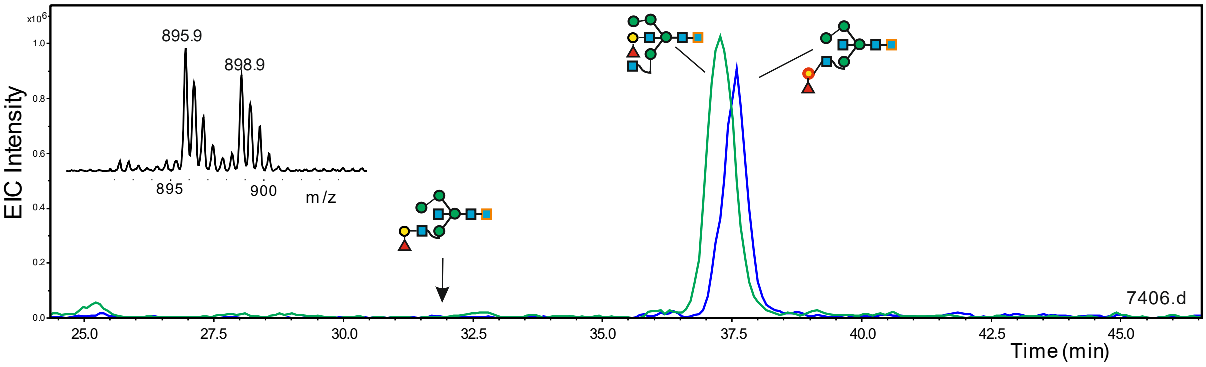

**Supplementary Fig. 9** Hybrid-type products of FucT-3. The blue EIC is the product obtained from the conversion of Man5GnF$^{6}$bi with b3GalT as shown in Fig. 4 (peak **a**). The green EIC shows - in the same analysis - the product obtained from brain peak 2, which obviously differs from the isomer elongated on the 3-arm.

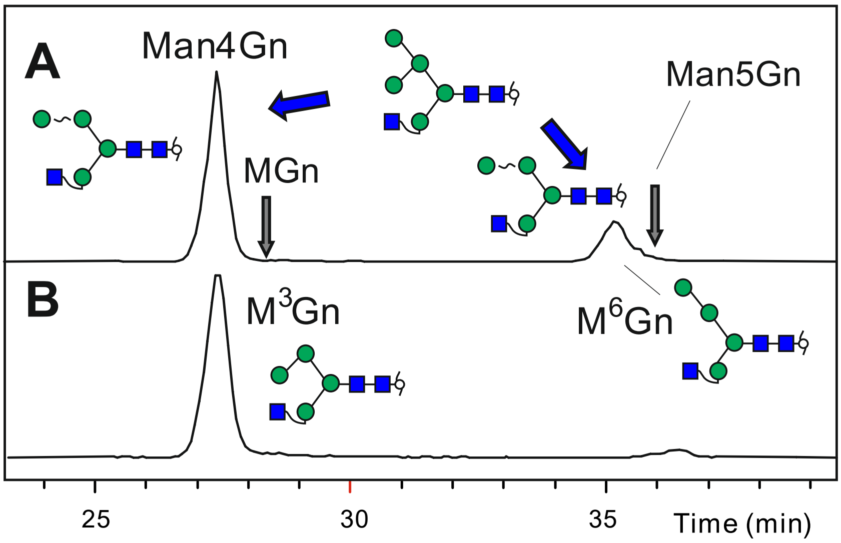

**Supplementary Fig. 10** Determination of the exact structure of Man4Gn. Panel **A** shows the result of a limited digest of Man5Gn with jack bean α-mannosidase. Panel **B** depicts the same EICs trace after treatment with an α1,6-specific mannosidase. Obviously, the large peak in panel **A** was the isomer with the upper arm mannose in 3-linkage. Removal of the 6-linked mannose causes a profound forward shift on the PGC column, which is considered useful for concluding on the mannose arrangement in other Man4 / Man5 families. The elution positions of the substrate Man5Gn and the final product MGn are indicated by arrows. The bias towards $M^3Gn$ is more pronounced than reported previously [7]

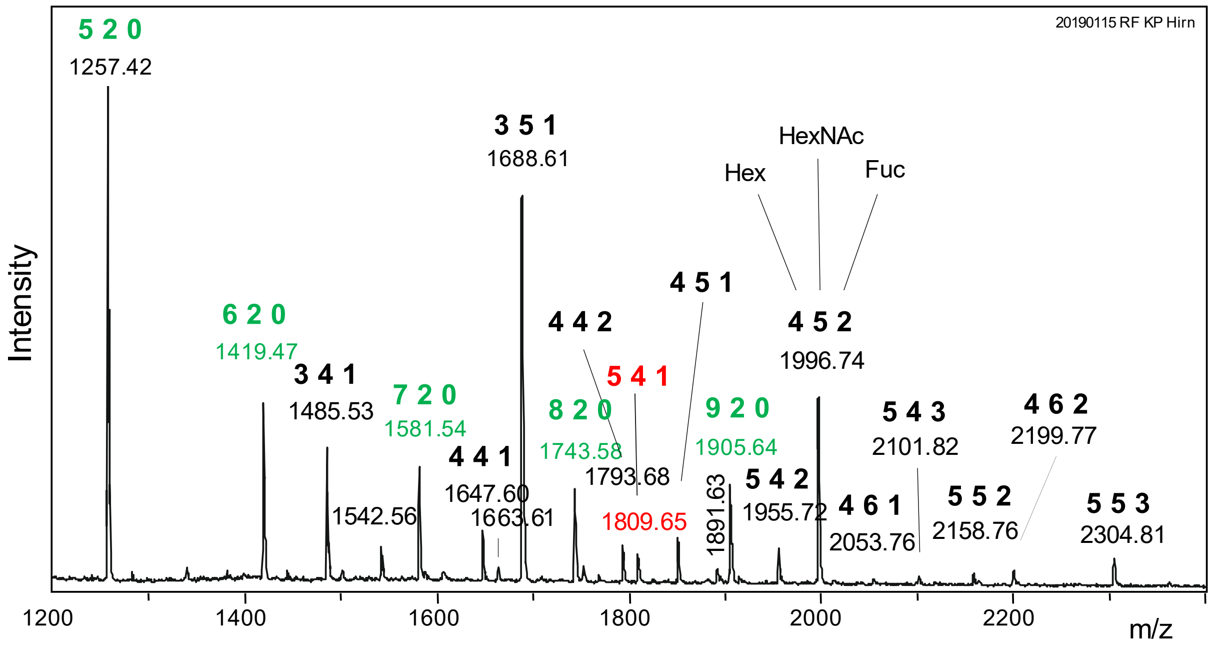

**Supplementary Fig. 11** MALDI -TOF MS of neutral porcine brain N-glycans. The peak for composition 5 Hex, 4 HexNAcs and 1 Fuc is emphasized in red.

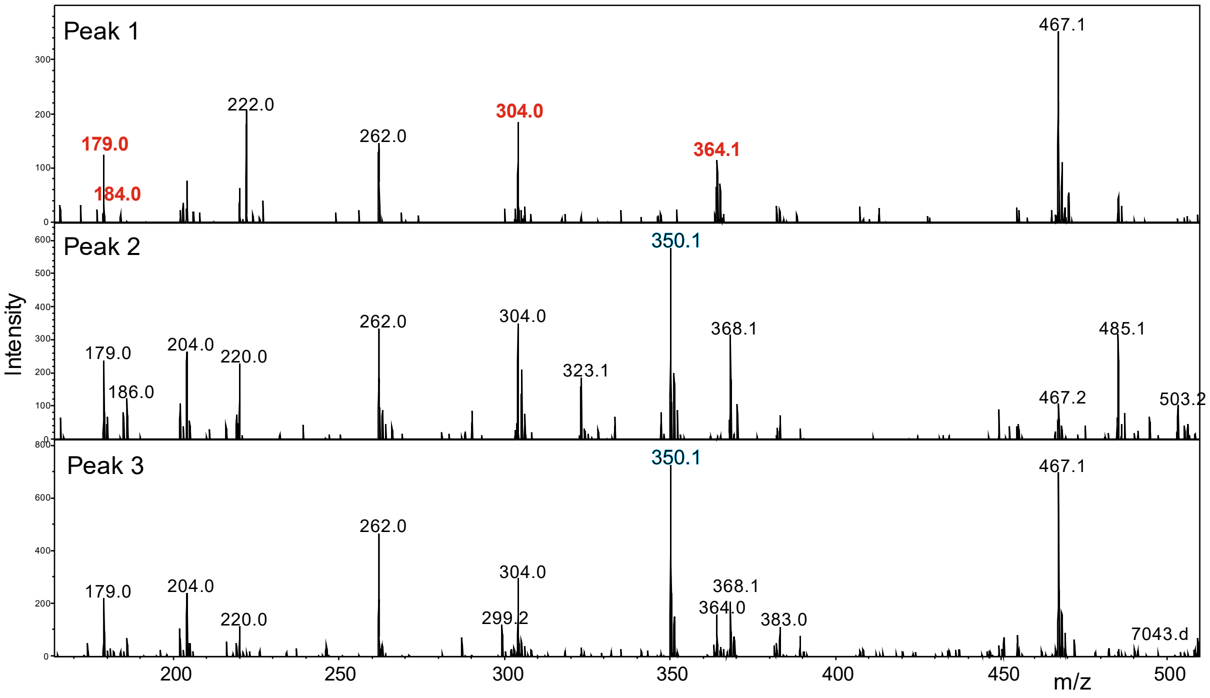

**Supplementary Fig. 12** Negative mode MS/MS of H5N4F1 brain peaks. The spectrum of peak 1 clearly indicates a LeX as opposed to a LeA structure [8]. The peaks relevant for distinction of LeX and LeA are printed in red. Peaks characteristic for core fucosylation are highlighted in blue.

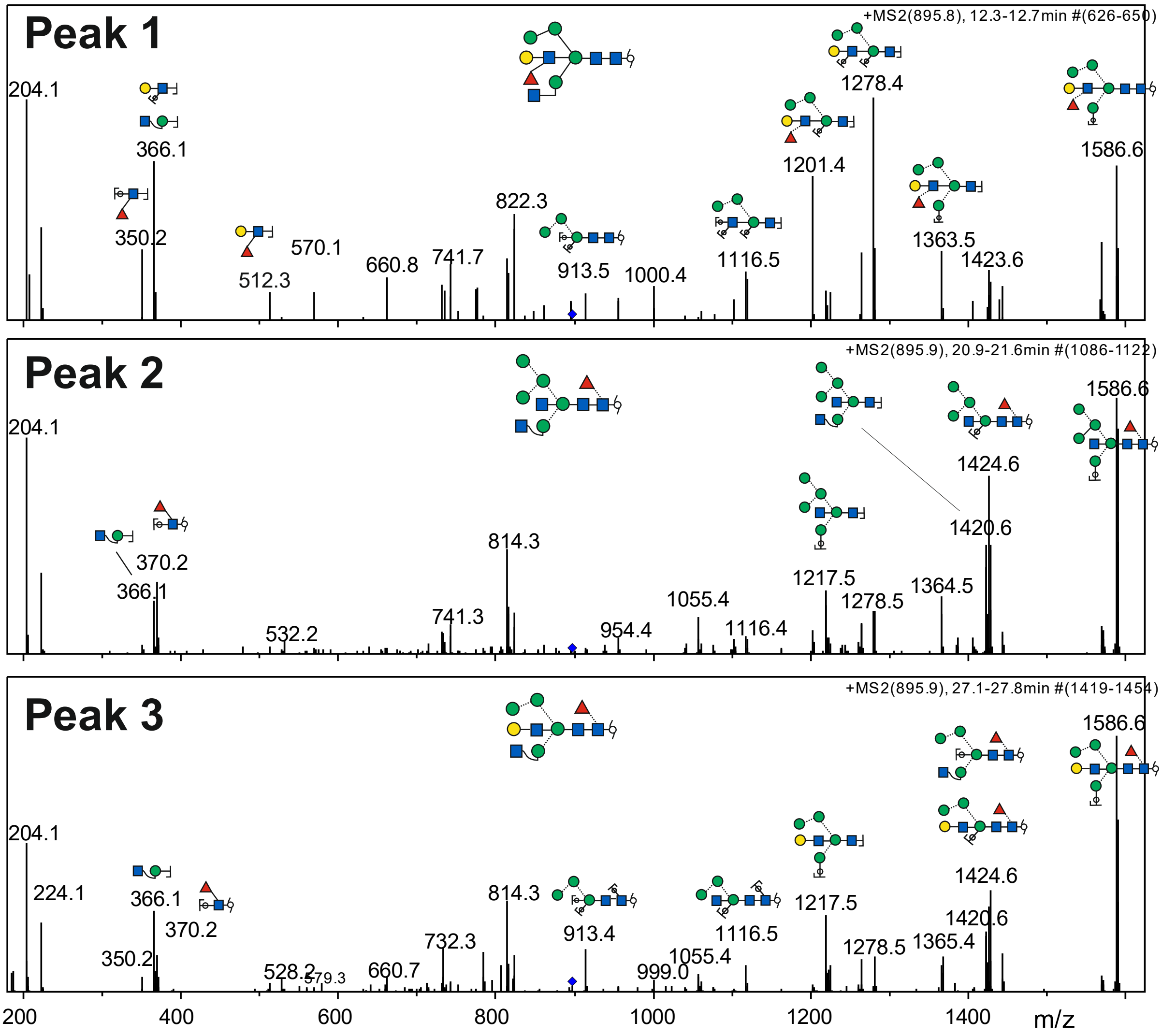

**Supplementary Fig. 13** Positive mode MS/MS of the hybrid-type H5N4F1 glycans of porcine brain. Fragments obtained from m/z = 895.9 precursors ($MH^+$ = 1789.67) of peak 1, 2 and 3 are shown in with tentative assignment of y and b fragment ions. Note the general dominance of the GlcNAc-loss fragment.

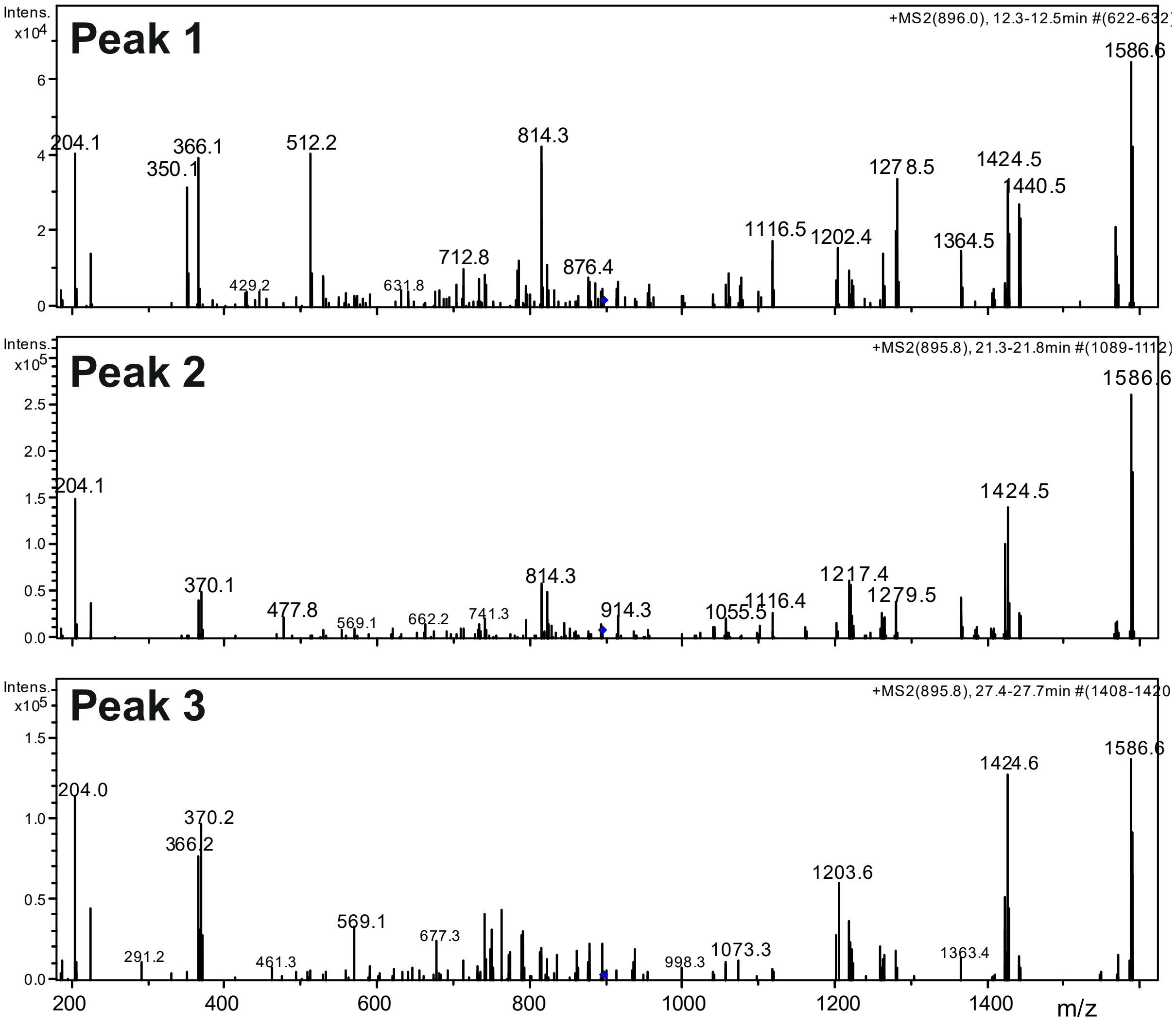

**Supplementary Fig. 14**: Positive mode MS/MS of the hybrid-type H5N4F1 glycans of human brain. Fragments obtained from m/z = 895.9 precursors ($MH^+$ = 1789.67) of peak 1, 2 and 3 are shown.

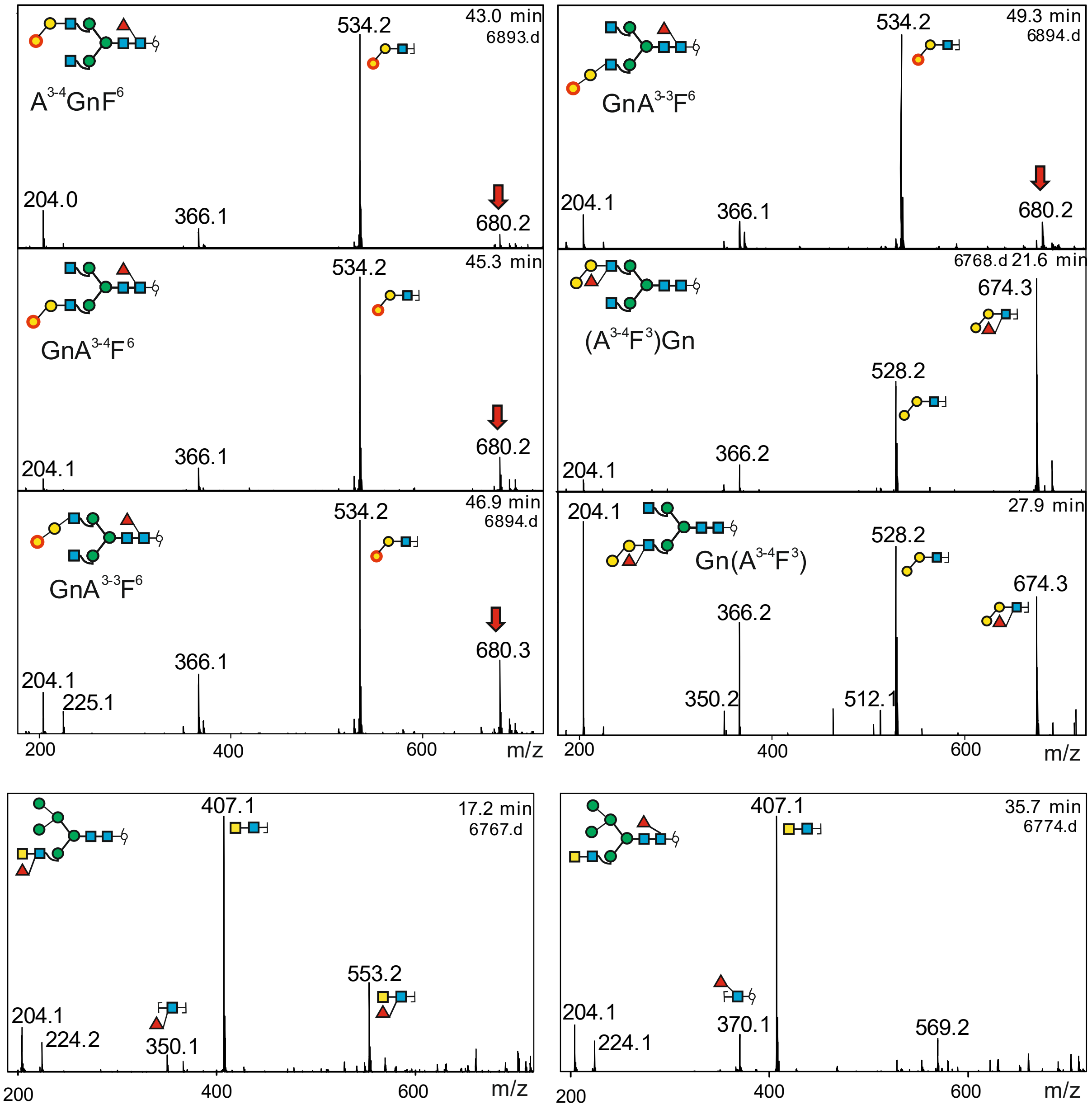

**Supplementary Fig. 15** Relevant sections of the positive mode MS/MS spectra of alfa-galactose and of GalNAc containing N-glycans with core- or LeX-fucose. The core-fucosylated structures are $^{13}C_6$-galactose labelled. Red arrows label fragments unquestionably resulting from re-arrangements.

##### Exposition of the N-glycan nomenclature system "proglycan" used in this work

In brief: "Proglycan" lists the terminal tips of an N-glycan in counterclockwise manner. The 6-arm, then 3-arm, then core fucose and finally bisection are indicated. The **terminus terms** are given such that a minimal number of characters nevertheless results in complete an unambiguous definition of the structure.

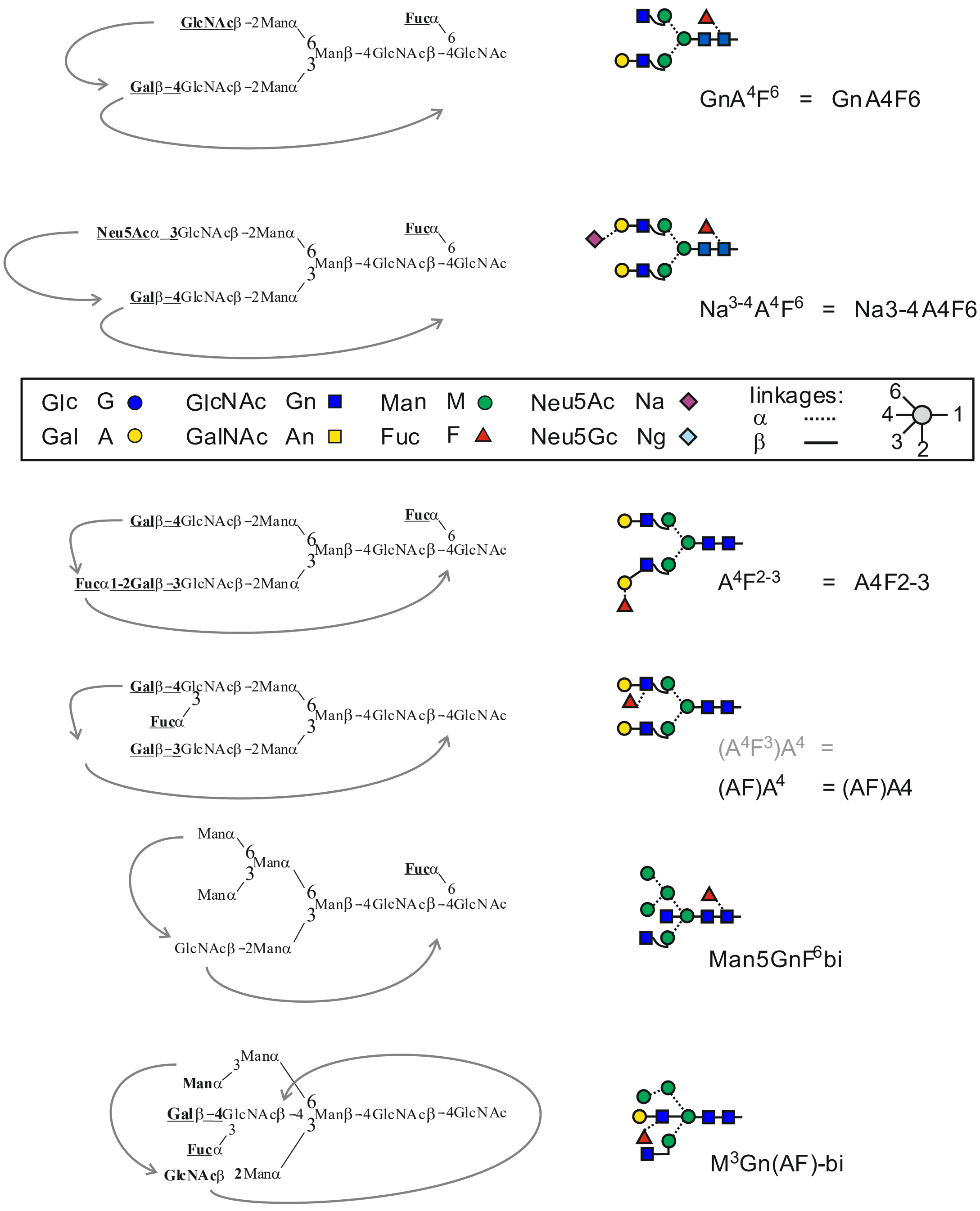

**Supplementary Fig. 16** Outline of the Proglycan nomenclature

The system starts with the term GnGn for the simple diantennary glycan with two terminal GlcNAc residues as coined by Harry Schachter [9] and also used by others [10]. Elongation of these antennae proceeds in a highly coordinated and defined manner. Thus, giving the very last residue of an antenna necessarily defines the following sequence. The residues are given by the one letter code, where A stands for galactose [11]. Modified monosaccharides such as N-acetylglucosamine are labelled by a second letter in small case, e.g. GlcNAc = Gn. In the case of sialic acids, Na and Ng stand for N-acetyl- or N-glycolyl-neuraminic acid, respectively (Supplementary Fig. 16).

The educated reader will immediately object that the situation is more complex as galactose is most often but not always found β1,4-linked to GlcNAc and sialic acids may be linked to the 3 or the 6 position of galactose. These ambiguities are eliminated by superscripts. $A^4$ stands for a Galβ1-4GlcNAcβ1-2Manα1-3 branch IF the **terminus term** $A^4$ occupies the second position as in the figure below.

Sialic acid is usually linked to galactose: $Na^3$-$A^4$ could thus be written for the terminus of the 6-arm in the figure below. But we rather don´t waste space for the self-evident. This **terminus term** $Na^3$-$A^4$ is written as $Na^{3\text{-}4}$. Fortunately, information is not lost together with formatting.

Fucosylation and bisecting GlcNAc are key issues of the current work. Core-fucosylation and bisection each have their own **terminus term** - F and bi.

The term F, however, is somewhat unclear in an environment dealing with insect and plant derived glycoproteins. Hence, it appears advisable to write $F^6$ for full definition of a mammalian type N-glycan, while $F^3$ denotes the core-1,3-fucose occurring in plants or insects.

Outer arm fucosylation occurs in three variants: the blood group H determinant Fucα1,2Gal, further Lewis A and Lewis X, where biosynthetic necessities always lead to branched structures. For the linear blood group H determinant, the IUPAC convention of connecting sugar by a line is adopted. Fucα(1-2)Galβ(1-3)…. becomes $F^2$-$A^3$. It appears equally clear and catchier to just write $F^{2\text{-}3}$. For branched structures, brackets are used, however, such that both branching sugars are placed within the bracket. The antenna Galβ-4(Fucα-3)GlcNAcβ … becomes ($A^4F^3$) – the GlcNAc is self-evident. The linkage symbols are also self-evident so that a LeX determinant becomes (AF), whereas a LeA structure would be (FA).

The hyphen comes back in the admittedly very special case of substituted bisecting GlcNAc. The "bi" is then extend to $A^4$-bi, or actually just A-bi as b3GalT does not act on the bisecting GlcNAc. Similarily, a LeX determinant is written as (AF)-bi to distinguish it from regular outer arm modifications.

We are fully aware that this system may appear unnecessary in the light of the numerous system used for decades [2], [12] or the highly sophisticated general annotation scheme for IT purposes [13]. We nevertheless consider the Proglycan system the first and only one that condenses N-glycan structures into a – in most cases – handy and easy to understand short string,

**List of possible terminus terms:**

in both first (6-arm) and second (3-arm) position:

| | |
|---|---|
| Gn | a GlcNAc linked to the invariant pentasaccharide core |
| U | indicates no substituent of the β-mannose. |
| A | Galactose linked to GlcNAc, linkage not specified |
| $A^4$ / A4 | Galactose β1,4-linked to GlcNAc |
| $A^3$ / A3 | Galactose β1,3-linked to GlcNAc |
| (AF) / ($A^4F^3$) / (A4F3) / Lx | Lewis X determinant |
| (FA) / ($F^4A^3$) / (F4A3) / La | Lewis A determinant |
| $F^{2\text{-}3}$ / $Lh^3$ | Blood group H determinant on type I chain |
| $F^{2\text{-}4}$ / $Lh^4$ | Blood group H determinant on type II chain |
| Na | Sialic acid linked to Gal-GlcNAc, linkages not specified |
| $Na^{6\text{-}4}$ / Na6-4 | Neu5Ac-α2,6-Gal-1,4-GlcNAc- |
| $Na^{3\text{-}3}$ / Na3-3 | Neu5Ac-α2,3-Gal-1,3-GlcNAc- |
| ($Na^6Na^{3\text{-}3}$) | Neu5Ac-α2,6(Neu5Ac-α2,3-Gal-1,3)-GlcNAc- (as in fetuin) |

Possible in first (6-arm) position:

| | |
|---|---|
| M | 6-arm mannose of the invariant pentasaccharide core |
| $M^3$ / M3 | a mannose in 3-linkage to 6-arm mannose of the core |
| $M^6$ / M6 | a mannose in 6-linkage to 6-arm mannose of the core; rare |
| [GnGn] | GlcNAc-β1,6-(GlcNAc-β1,2-)Man |
| [$A^4A^4$] | branched 6-arm with two Gal residues (in analogy to [GnGn]) |

Possible in second (3-arm) position:

| | |
|---|---|
| M | 3-arm mannose of the invariant pentasaccharide core |
| $M^2$ / M2 | a mannose in 2-linkage to the invariant pentasaccharide core |
| $M^{2\text{-}2}$ / M2-2 | two mannoses in series |
| [GnGn] | GlcNAc-b1,4-(GlcNAc-b1,2-)Man (in analogy to [GnGn]) |
| [$A^4A^4$] | branched 3-arm with two Gal residues (in analogy to [GnGn]) |

Possible in the extension positions – if present; fucosylation before bisection

| | |
|---|---|
| $F^6$ F | α1,6-fucosylation of reducing end GlcNAc |
| $F^3$ F | α1,3-fucosylation of reducing end GlcNAc |
| $F^3F^6$ | difucosylation of reducing end GlcNAc |
| X | β1,2-xylosylation of the β-mannose (written before fucosylation) |
| bi | indicates presence of a bisecting GlcNAc |
| A-bi | bisecting LacNAc |
| (AF)-bi | bisecting Lewis X |
